# Hydrophilic Polydopamine (hPDA) Fueled Bioglue Enhances Tissue Adhesion and Promotes Healing of Avascular Meniscus Tears

**DOI:** 10.64898/2026.07.14.738365

**Authors:** Hasan Rafsan Jani, Maya A. Jeremias, Aminah T. Sarowar, M. Nurul Islam, Chang H. Lee, Solaiman Tarafder

## Abstract

Avascular meniscus tears exhibit minimal intrinsic healing and often progress to joint degeneration due to restricted biological repair capacity and inadequate restoration of tissue-level structure and function. Here, we report a hydrophilic polydopamine (hPDA) fueled bioglue platform that overcomes the solubility limitations of conventional polydopamine (PDA) and enables functional repair of avascular meniscus injuries. Water-soluble hPDA was synthesized via controlled depolymerization and recrystallization, yielding monomeric and oligomeric species rich in catechol, amine, and hydroxyl functionalities. Incorporation of hPDA into fibrin bioglues markedly enhanced mechanical performance, producing 520–525% increases in lap-shear modulus, 165–190% increases in adhesive strength, and a 160% increase in compressive modulus relative to fibrin controls, while degradation was markedly attenuated over 14 days. hPDA exhibited excellent cytocompatibility in both 2D and 3D cultures. In a bovine avascular meniscus explant model, hPDA fueled bioglues promoted tissue integration and aligned collagen remodeling, restoring interfacial mechanics with a 488% increase in tensile modulus and up to 150% higher pull-out strength after 6 weeks. These findings establish hPDA as a versatile bioadhesive building block with strong potential for repairing avascular meniscus tears and other mechanically demanding connective tissues.

**Graphical Abstract:** 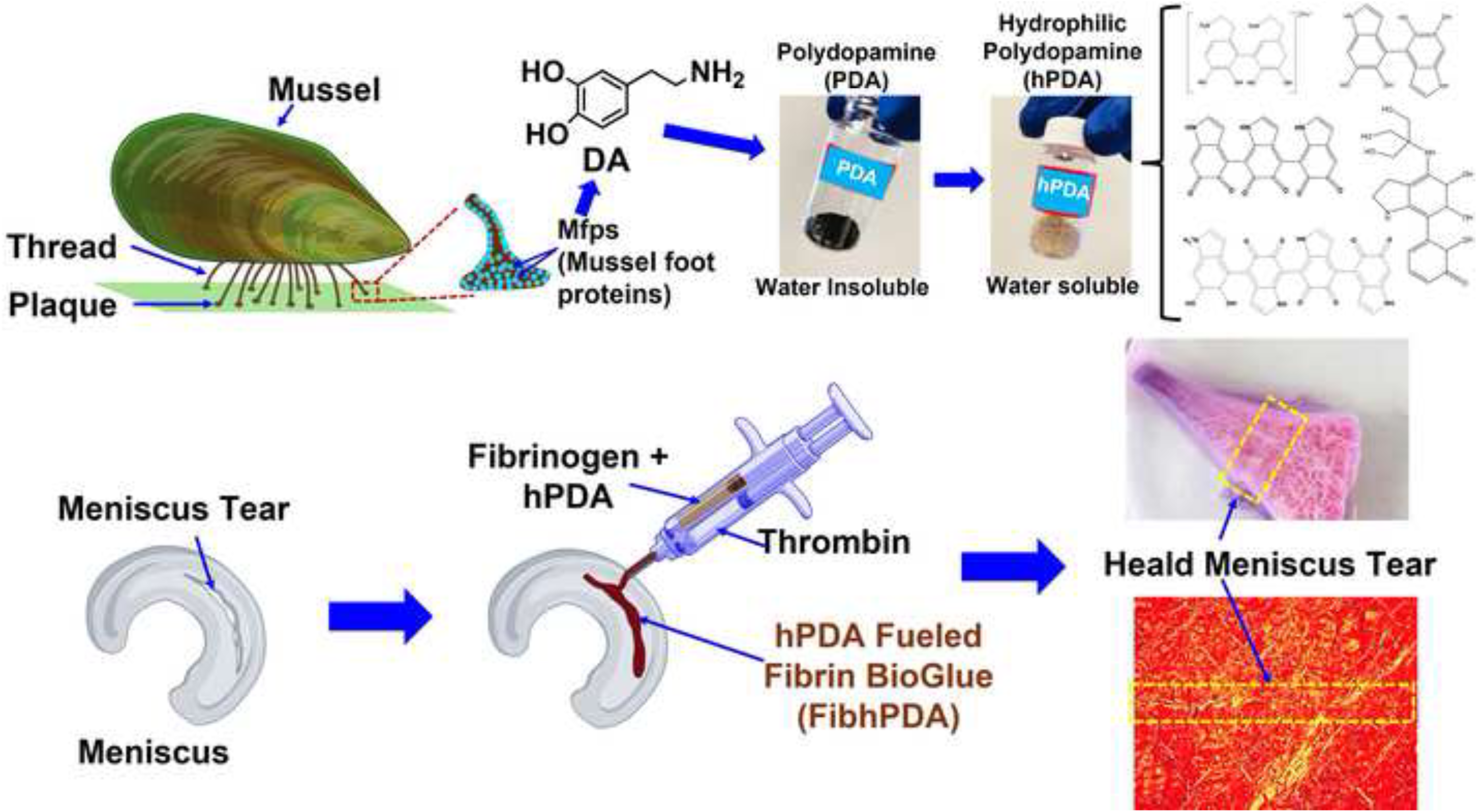

## 1. Introduction

The menisci of the knee are crucial for preserving joint alignment, providing shock absorption, and facilitating the distribution of forces across the joint area. Their intricate design and varied biochemical makeup are essential to their performance [1, 2]. The peripheral third of the meniscus is richly supplied with blood vessels and consists of a thick fibrous matrix that is home to cells resembling fibroblasts. On the other hand, the inner third is made up of avascular cartilaginous tissue, which is inhabited by cells similar to chondrocytes [3]. The central portion is identified as fibrocartilaginous and hosts a hybrid collection of both fibroblasts and chondrocytes. Each year in the U.S., about one million individuals undergo surgical procedures for the repair or removal of the meniscus [4, 5]. Suturing is an effective method for repairing tears in the vascularized outer third of the meniscus. However, tears within the inner avascular section do not naturally heal, frequently extending into the middle-third area and leading to reduced effective healing after repair. Such conditions often result in the spread of tears, deterioration of the meniscus, and ultimately, the development of osteoarthritis in the knee concerned [6, 7].

Meniscectomy, involving the partial or total removal of the meniscus, is a commonly executed surgery aimed at alleviating symptoms associated with meniscal injuries [8]. This approach, while initially effective, carries a significant long-term risk of osteoarthritis (OA) due to increased pressure on the joint surfaces [9]. Studies show that almost half of patients experience OA within 10 to 20 years following this intervention [6, 7, 8, 10, 11]. Following meniscectomy, allograft transplantation from deceased donors appears as a solution for mitigating long-term pressure problems. However, this approach is hampered by difficulties in finding suitable donors, potential size discrepancies, immune responses, disease transmission risk, graft shrinkage or extrusion, the possibility of transplant failure, and substantial financial burden [6, 7, 12].

Recent advances in tissue engineering and regenerative medicine hold the promise of overcoming the current limitations associated with existing meniscus repair and replacement strategies. Various surgical, bioengineering and regenerative strategies have been investigated to improve the healing of avascular meniscus injuries, including various biomaterials, stem/progenitor cells, bioglues and/or bioactive cues [13, 14, 15, 16, 17, 18, 19, 20]. In animal models like rats, sheep, and mini-pigs, delivering autologous mesenchymal stem/progenitor cells (MSCs) has demonstrated a moderate degree of effectiveness in promoting meniscus repair [21, 22, 23, 24]. A recent study demonstrated improved healing of meniscus tears through the use of modified cartilage progenitor cells (CPCs) with CXCR4, highlighting its potential as a promising therapeutic approach [25]. Researchers have further investigated the use of chemotactic cues and techniques to soften cell nuclei, aiming to promote the migration of internal cells and improve meniscus healing [26]. Alongside the exploration of cellular therapies, a growing body of research investigates various biomaterial grafts and tissue adhesives, aiming to promote healing in the avascular region of the meniscus [27, 18]. Despite significant research efforts, achieving complete functional healing of avascular meniscus tears, including the full restoration of their original biochemical composition and biomechanical properties, remains a challenge. This is particularly concerning given the widespread occurrence of meniscus injuries and their substantial impact on individuals’ quality of life. Consequently, a consistent and effective therapeutic strategy to promote healing in the inner meniscus is urgently needed.

Polydopamine (PDA) shares structural similarities with the sticky proteins used by marine mussels, allowing it to strongly adhere to various surfaces even in wet environments [28, 29, 30]. PDA possesses exceptional wet adhesion due to its active groups, adhering to diverse materials via covalent and noncovalent bonds [31]. PDA showcases remarkable performance in radical scavenging, ultraviolet shielding, photothermal conversion, electrochemical functionality, and biocompatibility. Given its versatility and straight forward synthesis from DA(3,4-dihydroxyphenethylamine), PDA finds extensive application across the biomedical, energy, catalyst, consumer, industrial, and military sectors, among others [30, 32, 33, 34, 35].

PDA exists as a thin film on different substrate surfaces, and it also forms nano to micro-sized spherical particles in solution [36]. Neither the thin film nor the particles dissolve in water, showing limited solubility in other organic solvents, which limits its application in many tissue engineering applications. Although PDA thin films and nano/microparticles are insoluble in most solvents, they are known to be degradable under highly basic conditions. Exposure of PDA films or particles to strong bases leads to their breakdown into smaller molecular fragments, primarily monomeric and oligomeric units containing two to four repeating structures. This degradation process is likely driven by the disruption of intermolecular hydrogen bonding, charge transfer interactions, and *π*–*π* stacking, which collectively contribute to the dissociation of PDA aggregates and render them water-soluble [37]. To date, there have been no reports describing the isolation or synthesis of these soluble PDA species. Here, we report for the first time a straightforward extraction method to obtain these soluble PDA fractions. Unlike conventional water insoluble PDA, our synthesized hydrophilic polydopamine (hPDA) exhibits excellent solubility and stability in aqueous environments. This breakthrough opens new opportunities for PDA-based materials, enabling a wide range of biomedical and tissue engineering applications that were previously inaccessible due to solubility limitations.

In this study, we developed and evaluated a hPDA fueled bioadhesive for meniscal tissue repair and regeneration. hPDA was synthesized through oxidative polymerization of DA followed by hydrophilic modification. FTIR confirmed retention of aromatic and amine groups with the appearance of new hydroxyl and carbonyl peaks, while mass spectrometry (MS) verified oligomeric and polymeric species formation with many functional groups available for strong tissue adhesion. SEM revealed a transition from the granular structure of PDA to a well-defined, plate-like morphology of hPDA. Incorporating hPDA into fibrin bioglue improved tissue adhesion, mechanical strength, viscoelasticity, and degradation control. Cytocompatibility assays confirmed excellent bioactivity of our synthesized hPDA. In a bovine meniscus explant model, hPDA fueled formulations enhanced tissue integration, collagen deposition, and interfacial strength compared to controls. Overall, this research establishes a promising bioadhesive platform capable of enhancing the healing of avascular meniscus tears. By improving tissue integration and mechanical restoration at the repair interface, this approach has the potential to support improved post injury joint function and long-term quality of life. Together, these findings represent a significant milestone in the translational development of hPDA fueled bioglue technologies for the functional repair of avascular meniscus tears and highlight their potential applicability to other mechanically demanding musculoskeletal connective tissues.

## 2. Results

### 2.1. Physicochemical Characterization of hPDA

SEM analysis revealed distinct morphological differences between PDA and hPDA (**Fig. 1a and 1b**). Oxidative polymerization of DA yielded PDA particles with sizes ranging from the nanoscale to the microscale. The synthesis of hPDA from PDA was achieved through controlled depolymerization in a basic medium, followed by recrystallization in excess ethanol and subsequent air drying (**Fig. S1**). The resulting hPDA displayed well-defined elongated, rod-like, and plate-like morphologies, distinct from the spherical structure of PDA. These morphological transformations were further confirmed by UV–Vis spectroscopy, where hPDA exhibited characteristic absorbance features markedly different from those of DA precursor (**Fig. 1c**). FTIR spectra of DA, PDA, and hPDA (**Fig. 1d**) illustrating the structural evolution and functional group retention during oxidative polymerization of DA to PDA and subsequent synthesis of hPDA from PDA.

**Figure 1:**
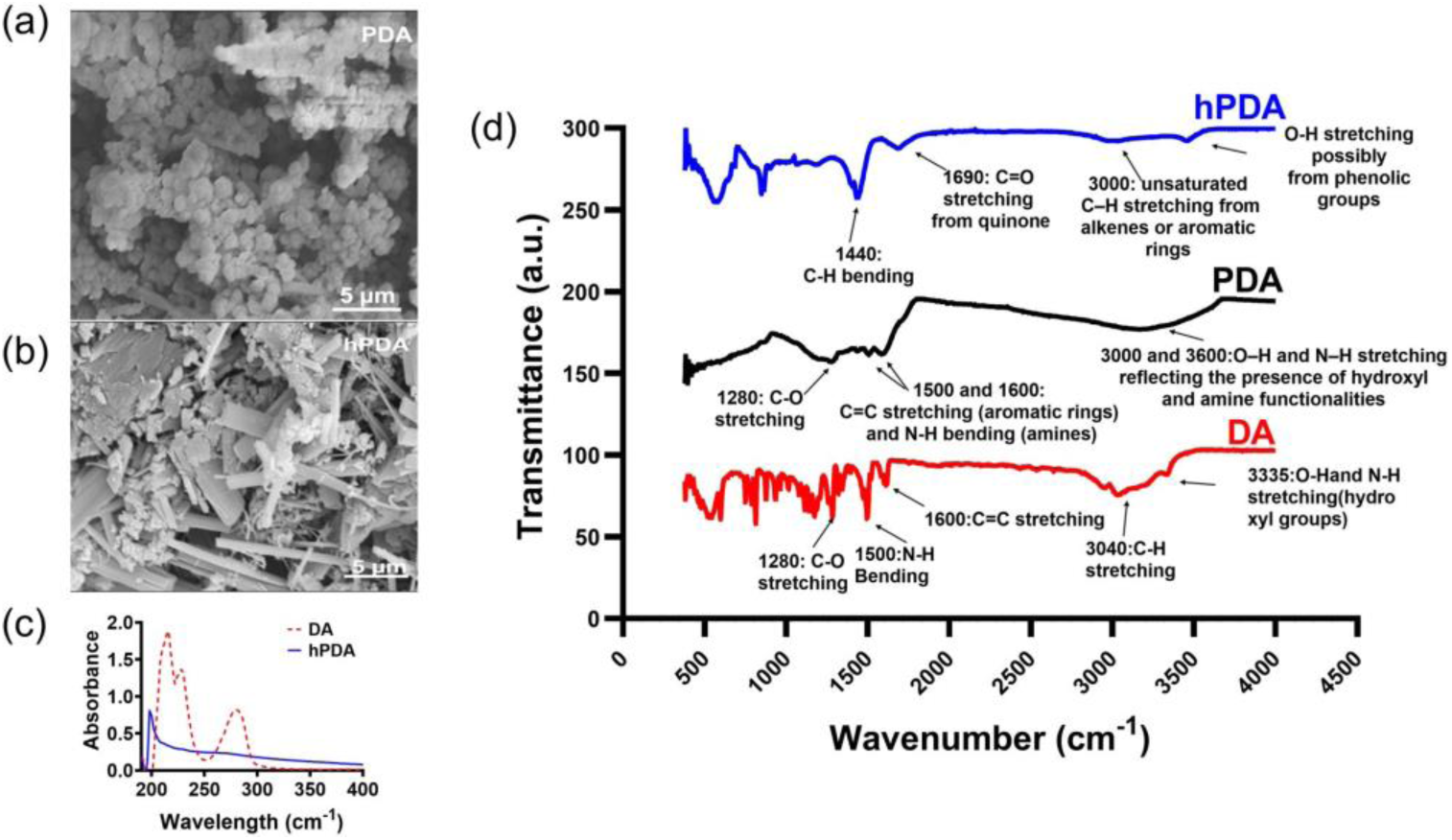
Structural, morphological, and spectroscopic characterization of DA, PDA, and hPDA. (a–b) Scanning electron microscopy (SEM) images of PDA and hPDA powders reveal distinct morphology evolution driven by synthesis and post-processing conditions. PDA forms aggregated particulate clusters, whereas hPDA exhibits plate-like crystalline morphologies consistent with solvent-mediated crystallization. (c) UV–Vis absorbance spectra illustrating the structural evolution from DA to hPDA. Dopamine shows characteristic sharp absorption bands in the 200–300 nm region, while hPDA exhibits a broad, featureless absorbance profile extending across the UV region. (d) FTIR transmittance spectra highlighting progressive chemical transformation from DA to PDA and hPDA, with retention of aromatic and amine functionalities alongside the emergence of hydroxyl and carbonyl groups, confirming successful synthesis of hydrophilic PDA species.

For DA (red spectrum), characteristic peaks are observed at 1600 cm^−1^, corresponding to aromatic C=C stretching vibrations, and at 1500 cm^−1^, attributed to N–H bending. A prominent peak near 1280 cm^−1^ is indicative of C–O stretching associated with phenolic groups. A broad absorption band centered around 3335 cm^−1^ is indicative to the O–H and N–H stretching vibrations, reflecting the presence of hydroxyl and amine functionalities. Additionally, a peak at 3040 cm^−1^ corresponds to C–H stretching vibrations from aromatic rings, while the peak at 2960 cm^−1^ is characteristic of aliphatic C–H stretching modes. Following oxidative polymerization to form PDA (black spectrum), the major functional group signatures are retained with noticeable peak broadening, reflecting the formation of a heterogeneous and cross-linked polymeric structure. The peak near 1590 cm^−1^ is assigned to aromatic and/or quinonoid C=C/C=N stretching, while the bands at 1500 cm^−1^ and 1280 cm^−1^ correspond to N–H bending and C–O stretching, respectively. A broad absorption band between 3000 cm^−1^ and 3600 cm^−1^ is attributed to the stretching vibrations of O–H and N–H groups. This broad feature also suggests the presence of hydrogen-bonded water molecules within the PDA structure. The FTIR spectrum of hPDA (blue) shows similar features to PDA, confirming the retention of the PDA structural backbone after hydrophilic modification. Two distinct broad bands are observed near 3000 cm^−1^ and 3650 cm^−1^. The 3000 cm^−1^ band may be attributed to unsaturated C–H stretching from alkenes or aromatic rings, while the 3650 cm^−1^ band is characteristic of free –OH stretching vibrations, indicating the presence of unbound hydroxyl groups. Additionally, slight shifts and changes in intensity are evident in the 1100 − 1300 cm^−1^ region. Notably, a strong and broad peak appears at 1690 cm^−1^, corresponding to the stretching vibration of carbonyl (C=O) groups from quinone structures, which is absent in the DA spectrum. These spectral changes collectively confirm the successful oxidative polymerization of DA to form PDA, as well as the effective subsequent synthesis of hPDA from PDA.

The mass spectrum of DA (**Fig. 2a**) exhibits a relatively simple profile, dominated by its molecular ion and characteristic fragmentation products, consistent with its low molecular weight and small molecular structure. Dopamine has a molecular weight of 153.18 Da, and the peak at m/z 154 Da corresponds to the protonated DA ion [M+H]=m/z+1. Fragmentation peaks at m/z 137 Da and 119 Da are likely due to the loss of –NH_2_ and – OH groups, respectively which are typical for catecholamine compounds under ionization conditions [38]. In the mass spectrum of PDA (**Fig. 2b**), the persistence of low m/z peaks (119, 137, and 154 Da) indicates the presence of residual DA monomer and fragment-related species. In contrast, the emergence of higher m/z peaks such as 297, 402, and 453 Da confirms the formation of oligomeric structures. Specifically, m/z 297 Da is attributed to DA dimers, while m/z 402 and 453 Da may correspond to higher-order oligomers or DA dimers associated with tris(hydroxymethyl)aminomethane (Tris) used during synthesis (**Fig. 2c**) [39, 40, 41]. The presence of these higher mass peaks supports the successful oxidative polymerization of DA into PDA. The mass spectrum of hPDA (**Fig. 2c**) reveals a broader and more complex pattern than that of PDA, reflecting the breakdown of complex intra- and intermolecular bonds during the synthesis of hPDA from its insoluble PDA precursor. Peaks at m/z 353, 381, 437, and 475 Da could be assigned to oligomeric species, including dimers, tris-associated dimers, and trimers [40, 41, 37]. Notably, m/z at 588 Da tentatively originated from a tetrameric species as shown in **Fig. 2c** [42]. These findings confirm the structural complexity and successful functional modification of PDA to form hPDA.

**Figure 2:**
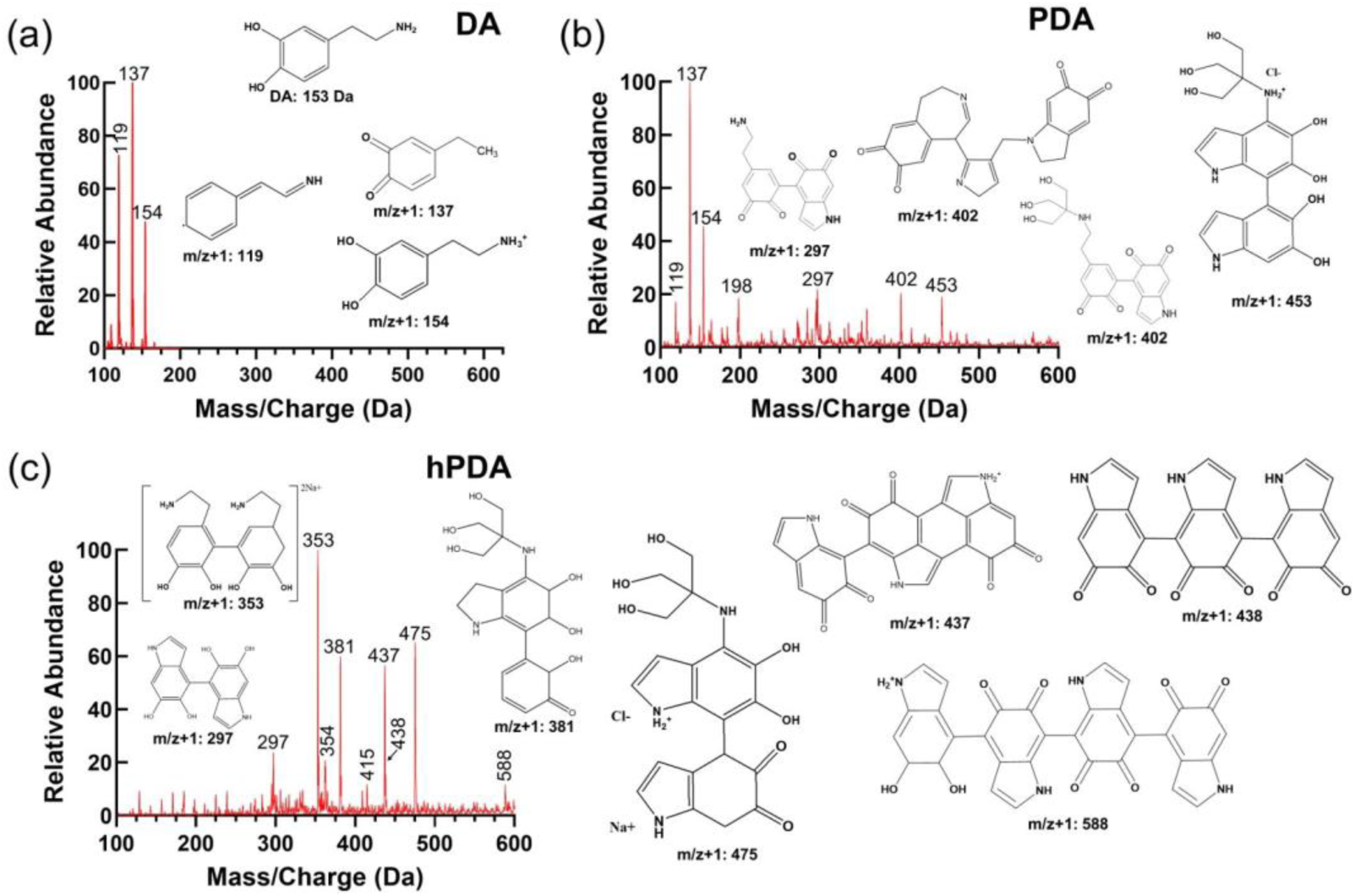
Mass spectrometric analysis of DA, PDA, and hPDA. (a) DA exhibits discrete low–molecular-weight ions corresponding to the monomeric species. (b) PDA displays a broad and heterogeneous mass distribution, reflecting the formation of higher-order oligomeric and polymeric species generated through oxidative polymerization. (c) hPDA reveals an enriched population of water-soluble oligomers with distinct sodium and chloride adducts, indicating successful depolymerization and chemical modification. Annotated peaks correspond to representative oligomeric structures consistent with the observed mass-to-charge (m/z) ratios. The presence of multiple functionalized oligomers highlights increased chemical accessibility and functional group availability, which underpin the enhanced interfacial reactivity and bioadhesive performance of hPDA.

### 2.2. Rheological Characterization

The rheological properties of the four hydrogel formulations revealed distinct flow behaviors and viscoelastic characteristics that were influenced by the incorporation of hPDA and genipin crosslinking. Viscosity measurements as a function of shear rate (**Fig. S2a**) demonstrated that all formulations exhibited shear-thinning behavior, with viscosity decreasing from approximately 2000-3000 Pa·s at low shear rates (0.1 s^−1^) to below 1 Pa·s at high shear rates (1000 s^−1^). The viscosity profiles largely overlapped across the tested shear rate range, indicating comparable injectability and flow characteristics among formulations. Strain sweep analysis (**Fig. S2b**) identified the linear viscoelastic region for each formulation, with the storage modulus (*G*′) remaining constant at low strains (0.1-10%) before decreasing at higher strains due to structural breakdown. All formulations maintained structural integrity up to approximately 100% strain, beyond which both *G*′ and loss modulus (*G*^′′^) values declined rapidly, indicating network disruption. Among the groups, FibhPDA exhibited the most robust mechanical response, maintaining higher modulus values over a broader strain range, suggesting enhanced network stability through hPDA incorporation.

Frequency sweep analysis (**Fig. S2c**) conducted within the linear viscoelastic region demonstrated gel-like behavior for all formulations, with *G*′ consistently exceeding *G*^′′^ across the frequency range of 0.1–100 Hz. The FibGenhPDA formulation exhibited the highest storage modulus, reaching approximately 3000–4000 Pa, followed by FibhPDA with intermediate values of approximately 1500–2000 Pa. In contrast, Fib and FibGen showed lower *G*′ values, with maxima near 1000 Pa. The relatively stable *G*^′′^ values across frequencies indicate good elastic recovery stability for all formulations.

### 2.3. Degradation Kinetics and Mechanical Performance of hPDA Fueled Bioglue

#### 2.3.1. Degradation Kinetics

The in vitro degradation behavior of the fibrin-based bioglues was systematically evaluated to assess the influence of hPDA incorporation and genipin crosslinking on their structural stability under physiological conditions (**Fig. 3a**). Fibrin (Fib) exhibited a rapid degradation profile, with complete mass loss by day 7, indicating its inherent susceptibility to enzymatic and hydrolytic breakdown. Incorporation hPDA into the fibrin matrix (FibhPDA) markedly slowed degradation, resulting in approximately 60% mass loss by day 14, suggesting that hPDA enhances network integrity through non-covalent intermolecular interactions and secondary crosslinking effects. In contrast, genipin-crosslinked fibrin formulations (FibGen and FibGenhPDA) showed substantially higher resistance to degradation due to covalent crosslinking between genipin and the primary amine groups of fibrin. Both FibGen and FibGenhPDA exhibited similar degradation profiles with minimal mass loss over 14 days, indicating that the genipin-mediated covalent crosslinking plays a dominant role in stabilizing the fibrin network. However, FibGenhPDA demonstrated slightly greater structural retention, highlighting a modest synergistic effect between genipin crosslinking and hPDA incorporation.

**Figure 3:**
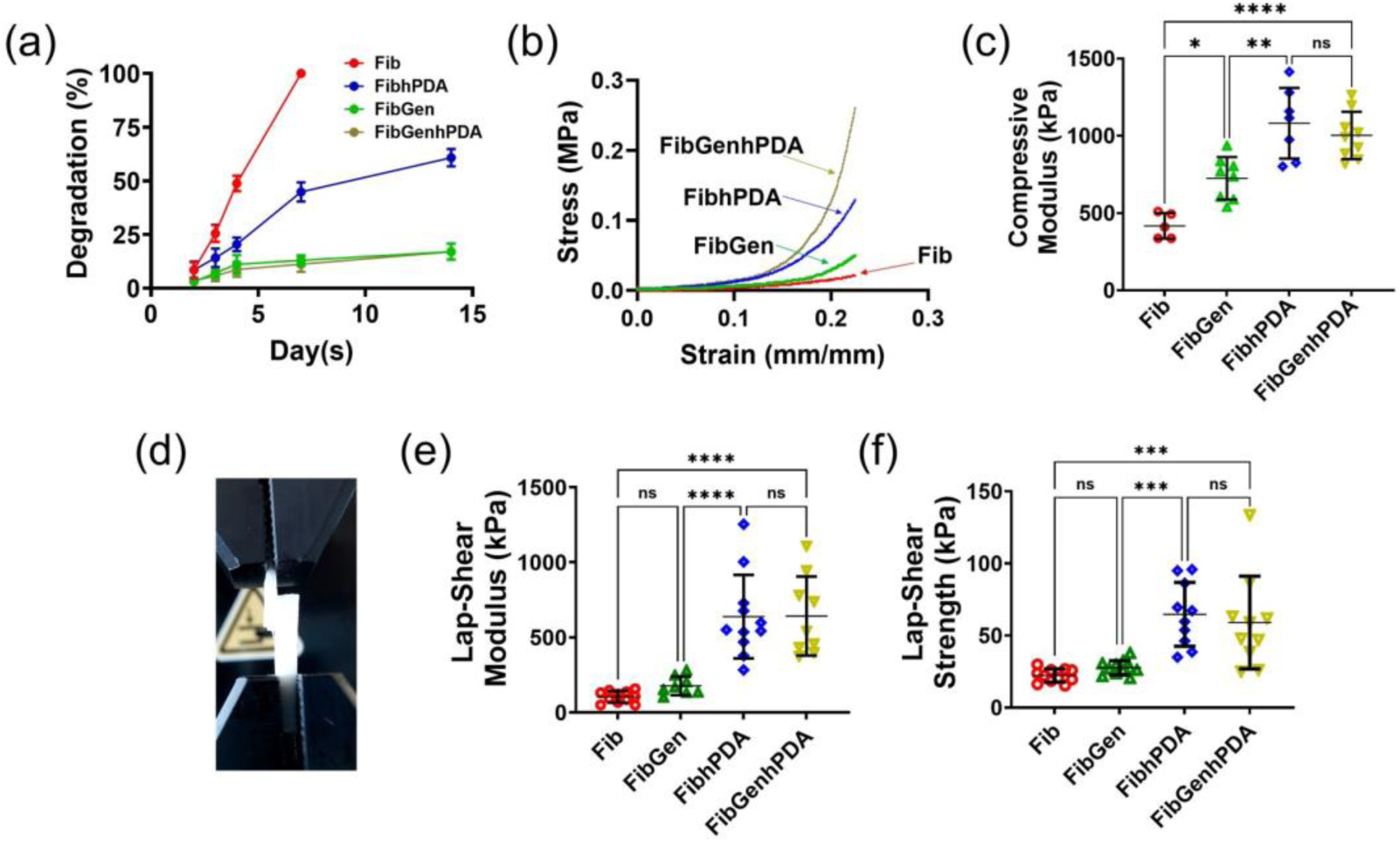
Degradation kinetics, compressive mechanics, and lap-shear interfacial adhesion test of hPDA fueled bioglues. (a) *In vitro* degradation profiles over 14 days demonstrate rapid mass loss for fibrin (Fib), moderated degradation upon hPDA incorporation (Fibh-PDA), and markedly attenuated degradation for genipin-crosslinked formulations (FibGen and FibGenhPDA), indicating enhanced structural stability. (b) Representative compressive stress–strain curves reveal characteristic nonlinear, J-shaped behavior for all formulations, with pronounced strain-stiffening in hPDA- and genipin-containing bioglues relative to Fib. (c) Quantitative analysis shows a stepwise increase in compressive modulus from Fib to FibGen, with further significant enhancement in FibhPDA and FibGenhPDA, reflecting synergistic reinforcement mechanisms (*n* ≥ 6). (d) Representative photograph of the lap-shear testing configuration using bovine meniscus strips, illustrating the defined bonded overlap area (5 × 10 mm) secured in custom grips and tested at a displacement rate of 0.1 mm s^−1^. (e) Lap-shear modulus measurements indicate substantial interfacial stiffening for hPDA-containing formulations ;(FibhPDA and FibGenhPDA) compared to Fib and FibGen (*n* ≥ 6). (f) Lap-shear strength follows the same hierarchical trend, with hPDA fueled bioglues achieving the highest adhesion values, while Fib and FibGen remain comparatively low (*n* ≥ 6). Statistical significance: ns, not significant; ^∗^ *p <* 0.05; ^∗∗^ *p <* 0.01; ^∗∗∗^ *p* ≤ 0.001; ^∗∗∗∗^ *p* ≤ 0.0001.

#### 2.3.2. Compression Test

The compression test results of the four bioglue formulations revealed significant differences in mechanical behavior, primarily influenced by the incorporation of hPDA and genipin. All formulations (**Fig. 3b**) exhibited characteristic nonlinear, J-shaped compressive stress–strain curves typical of soft hydrogel networks, showing an initial low-modulus region followed by pronounced stiffening at higher strains due to polymer network alignment and water expulsion. Among the groups, fibrin (Fib) demonstrated the lowest resistance to deformation, while genipin-crosslinked fibrin (FibGen) showed enhanced compressive strength. Incorporation of hPDA (FibhPDA and FibGenhPDA) further increased stiffness, producing a steeper elastic slope and improved load-bearing capacity across the tested strain range. Quantitative analysis of the compressive modulus (**Fig. 3c**) confirmed these trends. Fib exhibited the lowest modulus (420±82 kPa), consistent with its limited structural integrity. Genipin crosslinking (FibGen) significantly increased stiffness (725±137 kPa, p < 0.05 vs. Fib), attributed to covalent bond formation between genipin and fibrin amine groups. The inclusion of hPDA resulted in a further increase in stiffness for FibhPDA (1081±228 kPa, p < 0.0001 vs. Fib, p < 0.01 vs. FibGen), suggesting substantial reinforcement through non-covalent intermolecular interactions and secondary crosslinking effects. Although FibGenhPDA (1003±152 kPa) exhibited comparable stiffness to FibhPDA, the difference was not statistically significant, indicating that the reinforcing mechanisms of genipin and hPDA may partially overlap, yielding limited additive stiffening when combined.

#### 2.3.3. Lap-Shear Test

The lap-shear test results of the four bioglue formulations revealed significant differences in interfacial mechanical performance, primarily governed by the incorporation of hPDA rather than genipin alone. Representative lap-shear testing on bovine meniscus strips confirmed stable adhesive bonding across all groups, with failure occurring at the bonded interface (**Fig. 3d**). Among the formulations, fibrin (Fib) exhibited the weakest interfacial resistance, while genipin-crosslinked fibrin (FibGen) showed only modest improvement. In contrast, incorporation of hPDA (FibhPDA and FibGenh-PDA) resulted in a pronounced enhancement in both interfacial stiffness and load-bearing capacity. Quantitative analysis of lap-shear modulus (**Fig. 3e**) supported these observations. Fib exhibited the lowest modulus values (103±38 kPa), reflecting limited interfacial cohesion. FibGen showed a slight increase (176±71 kPa), corresponding to an approximate 71% increase relative to Fib; however, this difference was not statistically significant, indicating that genipin crosslinking alone had limited impact on adhesive stiffness. In contrast, inclusion of hPDA led to a substantial increase in modulus for both FibhPDA (637±278 kPa) and FibGenhPDA (642±263 kPa), representing an approximately 520–525% increase compared to Fib and a 260– 265% increase compared to FibGen. The absence of a significant difference between FibhPDA and FibGenhPDA suggests that hPDA provides the dominant contribution to interfacial reinforcement, with limited additive benefit from genipin when combined. A similar trend was observed for lap-shear strength (**Fig. 3f**). Fib (22±4.6 kPa) and FibGen (27±4.9 kPa) exhibited low adhesive strengths, with FibGen showing only a 23% increase relative to Fib and no significant difference between the two groups. In contrast, incorporation of hPDA significantly enhanced adhesion strength to 64±22 kPa for FibhPDA, corresponding to a 190% increase compared to Fib. FibGenhPDA achieved comparable values (58±32 kPa), representing a 165% increase relative to Fib and 110% increase relative to FibGen, with no statistically significant difference between the two hPDA-containing formulations. Collectively, these results demonstrate that hPDA plays a critical role in enhancing both interfacial stiffness and adhesive strength of the bioglue, likely through strong non-covalent interactions that promote effective load transfer and interfacial cohesion at the tissue–bioglue interface.

### 2.4. Cytotoxicity

In the 2D assay (**Fig. 4a**), hBMSCs exposed to hPDA at 0.5 and 1.0 mg/mL maintained normal spindle-shaped morphology and dense growth from Day 1 to Day 7. Cells showed predominantly green fluorescence, indicating high viability. Quantitative analysis confirmed that cell viability in both hPDA groups remained comparable to untreated controls throughout the study. In contrast, dopamine-treated groups at equivalent concentrations showed extensive red fluorescence and rapid cell loss beginning at Day 1, with viability decreasing to approximately 70% and 45% by Day 1 and falling below ∼20% and ∼10% by Day 2. By Days 3 and 7, dopamine-treated cultures exhibited near-complete cell death. The 3D assay in Fib and FibhPDA (**Fig. 4b**) supported these findings. hBMSCs encapsulated in FibhPDA constructs (6 mg/mL hPDA) remained highly viable from Day 1 to Day 7. Live/dead staining showed uniform green fluorescence with minimal red signal in both groups. Encapsulated cells gradually changed from rounded to elongated shapes by days 3–7, indicating active cell spreading. Quantitative analysis showed no difference in viability between Fib and FibhPDA groups, with live cell percentages consistently above 90%. Together, these results show that conversion of DA into hPDA eliminates dopamine-associated cytotoxicity and yields a highly biocompatible material suitable for direct cell-contact applications such as meniscal repair.

**Figure 4:**
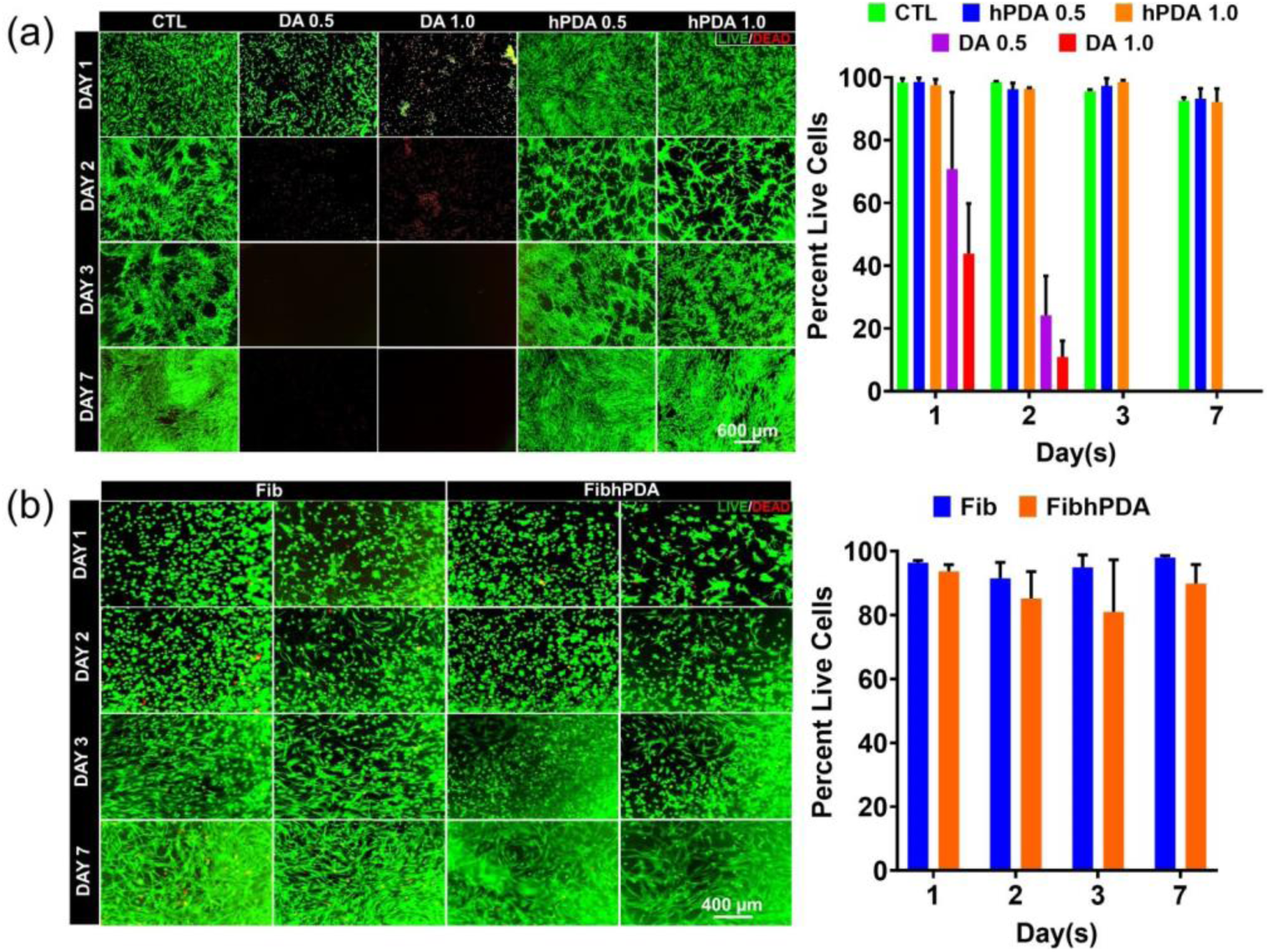
Cytocompatibility assessment of hPDA in two-dimensional (2D) and three-dimensional (3D) hBMSC cultures. (a) Representative live/dead fluorescence micrographs and corresponding quantitative viability analysis (bar graph) of hBMSCs cultured in 2D and exposed to DA (0.5 and 1.0 mg mL^−1^), hPDA (0.5 and 1.0 mg mL^−1^), or untreated control over 7 days. hPDA-treated cells retained normal spindle-shaped morphology and high viability comparable to controls across all time points, whereas DA exposure resulted in rapid and concentration-dependent cytotoxicity, with extensive cell death evident by day 2. (b) Live/dead staining of hBMSCs encapsulated within 3D fibrin and hPDA fueled fibrin bioglue (Fib vs. FibhPDA) demonstrates uniformly high cell viability from day 1 through day 7. Quantitative analysis confirms no statistically significant difference in viability between Fib and FibhPDA constructs at any time point, indicating that incorporation of hPDA does not compromise cell survival in a 3D microenvironment.

### 2.5. Explant Model for Avascular Meniscus Healing

Histological and collagen-specific staining showed clear differences in tissue integration and extracellular matrix organization at the meniscus repair interface, strongly influenced by hPDA incorporation (**Fig. 5a**). Hematoxylin and eosin (H&E) staining (**Fig. 5b**, top row) revealed poor interfacial continuity in the fibrin-only (Fib) group, with large gaps at the repair site consistent with rapid fibrin degradation and limited tissue bridging. The FibGen group showed partial defect filling with cell presence, but the regenerated tissue remained loosely organized and poorly integrated with native meniscus tissue. In contrast, hPDA fueled bioglues (FibhPDA and FibGenhPDA) exhibited tight apposition and continuous tissue across the interface, indicating improved adhesion and stable tissue bonding. Picrosirius Red staining (**Fig. 5b**, middle row) highlighted sparse and discontinuous collagen fibers in Fib and FibGen groups. In comparison, FibhPDA and FibGenhPDA showed dense, continuous collagen fibers spanning the repair region, suggesting enhanced collagen deposition and matrix remodeling. Under polarized light (**Fig. 5b**, bottom row), Fib and FibGen displayed weak birefringence and disorganized collagen, whereas hPDA fueled groups exhibited strong birefringence and well aligned fibers, most pronounced in FibhPDA, consistent with advanced collagen maturation.

**Figure 5:**
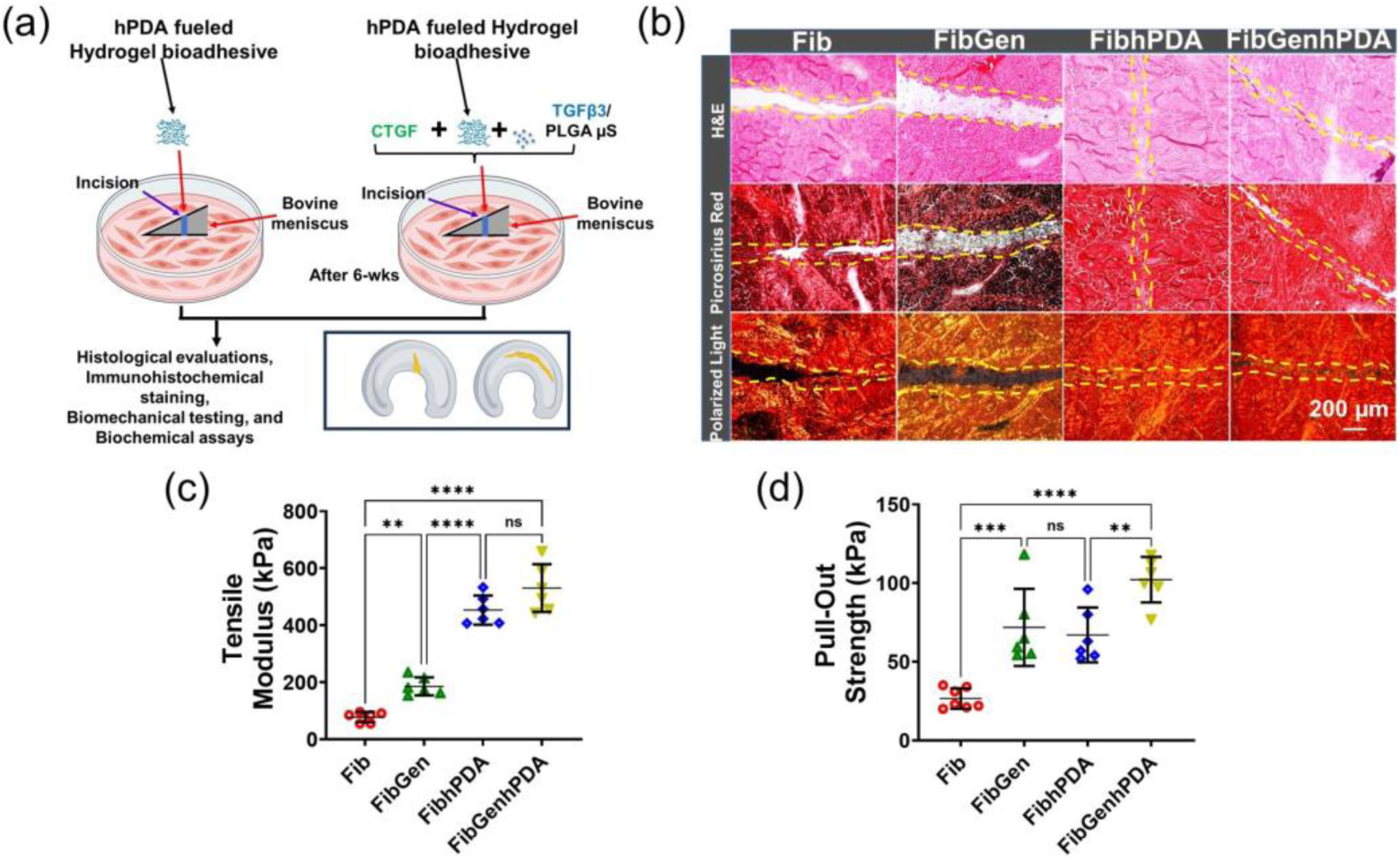
hPDA fueled fibrin bioglues promote functional integration and mechanical restoration in an avascular bovine meniscus explant model. (a) Schematic of the avascular meniscus explant repair model used to evaluate optimized hPDA fueled bioglues. Full-thickness inner-zone defects were filled with or without hPDA fueled fibrin bioglue supplemented with connective tissue growth factor (CTGF) and transforming growth factor-*β*3 (TGF-*β*3)-encapsulated PLGA microspheres. Treated explants were cultured for 6 weeks atop confluent (80%) human bone marrow–derived mesenchymal stem cell (hBMSC) monolayers, followed by histological, biochemical, and biomechanical analyses. (b) Representative histological sections after 6 weeks of culture stained with H&E, Picrosirius Red (brightfield), and Picrosirius Red under polarized light. Fib-treated explants exhibit poor interfacial continuity characterized by prominent gaps at the repair site, while FibGen shows partial defect filling with cell-populated regions. In contrast, FibhPDA and FibGenhPDA treatments demonstrate near-continuous tissue bridging, enhanced collagen deposition, and improved fiber alignment across the repair interface. Yellow dashed lines denote the original defect boundary. (c) Tensile modulus of repaired explants increases progressively from Fib to FibGen, with a marked enhancement observed in hPDA fueled formulations (FibhPDA and FibGenhPDA), indicating improved interfacial stiffness and load transfer. (d) Pull-out strength measurements reveal significantly enhanced interfacial bonding with hPDA incorporation, with FibGenhPDA achieving the highest resistance to interfacial failure. Data are presented as mean ± SD (*n* ≥ 6). Statistical significance: ns, not significant; ^∗^ *p <* 0.05; ^∗∗^ *p <* 0.01; ^∗∗∗^ *p <* 0.001; ^∗∗∗∗^ *p <* 0.0001.

### 2.6. Tensile Testing for Tissue Integration

Uniaxial tensile pull-out testing after 6 weeks explant culture revealed strong, formulation-dependent differences in mechanical integration at the repaired meniscus interface. Fibrin only (Fib) repairs exhibited the lowest tensile modulus (**Fig. 5c**), indicating poor load transfer. Genipin cross linking (FibGen) moderately increased both modulus (139% increase relative to Fib) and strength (170% increase vs. Fib), but incorporation of hPDA produced substantially greater improvements. FibhPDA showed a large increase in tensile modulus (482% increase vs. Fib), while FibGenhPDA achieved the highest stiffness overall (582% increase vs. Fib), indicating that hPDA is the primary contributor to interfacial load transfer, with genipin providing limited additional reinforcement. Tensile pull-out strength measurements (**Fig. 5d**) followed a similar but not identical hierarchy, highlighting complementary roles of hPDA and genipin in resisting interfacial failure. Fib failed at low stresses (26.57 ± 6.50 kPa), while FibGen increased strength to 71.80 ± 24.54 kPa (170% increase vs. Fib). FibhPDA achieved comparable strength (67.00 ± 17.44 kPa; 152% increase vs. Fib), whereas FibGenhPDA exhibited the highest pull-out strength (102.17 ± 14.47 kPa; 285% increase vs. Fib). Together, these findings indicate that hPDA primarily enhances tensile load transfer, while genipin contributes to peak interfacial strength, with their combination yielding the most robust functional integration of the healed meniscus interface.

### 2.7. Modulus Mapping By Nanoindentation

The effective elastic modulus (*E_Eff_*) mapping revealed pronounced differences in local micro-mechanical properties across the healing interface of meniscus explants (**Fig. 6a**), largely governed by the presence of hPDA within the bioglue formulations. High-resolution spatial mapping performed across the repair region demonstrated distinct mechanical discontinuities in non-hPDA groups, while hPDA-containing groups exhibited improved mechanical continuity and stiffness. In the fibrin-only (Fib) group, a marked reduction in *E_Eff_* was consistently observed within the interfacial region (highlighted by dashed red boxes), indicating poor tissue integration and the presence structural discontinuity. Although there was no structural discontinuity observed in FibGen group, the interfacial *E_Eff_* values remained significantly lower than those of the surrounding native tissue, reflecting incomplete mechanical restoration. In contrast, incorporation of hPDA (FibhPDA and FibGenhPDA) resulted in substantially more uniform modulus distributions across the healing interface, with elevated *E_Eff_* values that closely approached those of native meniscus tissue. These groups exhibited reduced spatial variability and improved stiffness continuity across the repair region, suggesting enhanced matrix deposition and integration at the microscale. Quantitative analysis of average *E_Eff_* values (**Fig. 6b**) confirmed these trends, showing significantly higher moduli in both hPDA-containing groups compared to Fib and FibGen.

**Figure 6:**
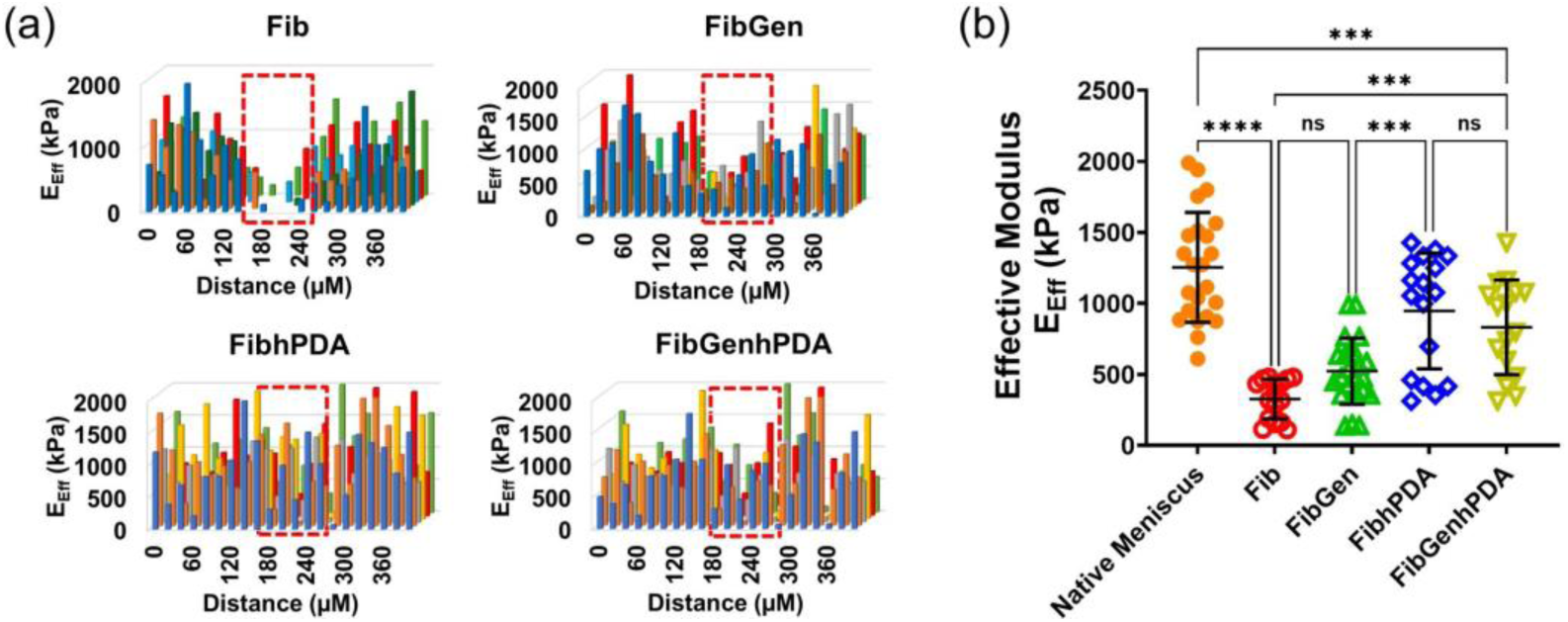
hPDA fueled bioglue enhances interfacial load-bearing and mechanical integration in avascular meniscus repair. (a) Spatial mapping of effective elastic modulus (*E*_Eff_) across the repair interface of bovine avascular meniscus explants after 6 weeks of culture, obtained by nanoindentation at 20 µm intervals. Fib-treated explants exhibit a pronounced low-modulus valley centered at the defect site (red dashed boxes), indicative of mechanical discontinuity and poor tissue integration. FibGen shows a narrower but persistent modulus depression at the interface. In contrast, hPDA fueled formulations (FibhPDA and FibGenhPDA) display uniform, plateau-like modulus profiles across the interface, reflecting continuous load-bearing matrix formation and improved mechanical coupling between adjacent tissue regions. (b) Quantitative comparison of average *E*_Eff_ values reveals that native meniscus exhibits the highest stiffness, while Fib-treated explants show a significant reduction in modulus relative to native tissue. FibGen demonstrates a modest increase that is not statistically different from Fib. Both hPDA fueled groups exhibit a substantial elevation in *E*_Eff_ compared to Fib and FibGen, approaching native tissue stiffness, with no significant difference between FibhPDA and FibGenhPDA. Data are presented as mean ± SD. Statistical significance: ns, not significant; ^∗^ *p <* 0.05; ^∗∗^ *p <* 0.01; ^∗∗∗^ *p <* 0.001; ^∗∗∗∗^ *p <* 0.0001.

## 3. Discussion

The inability of avascular meniscus tears to heal remains a central translational challenge in orthopaedic repair, arising from a convergence of biological constraints, poor interfacial adhesion, and inability to restore mechanical continuity under physiological loading. [6, 7, 8, 10, 11, 9, 43] Recent advances in meniscus biology and diagnostics have clarified that durable repair requires strategies that stabilize the repair interface early while enabling subsequent cell mediated remodeling and matrix maturation, rather than relying on defect filling alone. [9] In this context, the present study introduces hydrophilic polydopamine (hPDA) as a materials driven solution that directly addresses these interfacial and mechanical bottlenecks through strong adhesion, controlled degradation, and biological compatibility.

Catechol mediated adhesion, inspired by mussel foot proteins, provides a foundational paradigm for achieving robust bonding in wet and dynamic environments. [29] Polydopamine extended this concept to synthetic systems, offering substrate independent adhesion and broad chemical versatility. [28, 30, 31, 33, 44] However, the widespread translational use of polydopamine (PDA) in injectable or cell laden biomaterials has been limited by its intrinsic water insolubility, aggregation, and restricted molecular accessibility [36, 37]. In this work, controlled depolymerization and recrystallization (**Fig. S1**) yielded water soluble hPDA oligomers that preserve catechol rich chemistry while markedly enhancing functional group accessibility. Structural and spectroscopic analyses (**Fig. 1**) together with mass spectrometry (**Fig. 2**) confirmed retention of hydroxyl, amine, and carbonyl functionalities, which are known to mediate hydrogen bonding, metal coordination, and covalent coupling with collagenous matrices.[28, 30, 31, 44]

When integrated into fibrin bioglue, hPDA produced substantial improvements in viscoelastic stability (**Fig. S2**), degradation resistance (**Fig. 3A**), and compressive stiffness (**Fig. 3B and C**) without compromising shear thinning behavior required for surgical handling. Although genipin crosslinking is widely used to stabilize protein based hydrogels through covalent amine coupling [45], our data demonstrate that genipin alone provides only modest improvements in interfacial adhesion. In contrast, hPDA drove dominant increases in lap shear modulus and strength (**Fig. 3E and F**), highlighting that interfacial chemistry rather than bulk crosslink density is the primary determinant of functional tissue bonding. [31] The limited additive effect between genipin and hPDA suggests partially overlapping reinforcement mechanisms and is consistent with prior observations that increased bulk stiffness does not necessarily translate to improved tissue material integration. [27, 18] These findings align with emerging interface driven design principles in regenerative biomaterials.

Biological compatibility is a critical translational requirement for adhesive systems intended for load bearing soft tissues. While free dopamine exhibited severe cytotoxicity (**Fig. 4A**), consistent with its oxidative reactivity, [36, 33, 39] polymerization followed by transformation into hPDA fully mitigated these effects. hPDA supported sustained viability of hBM-SCs in both 2D and 3D environments (**Fig. 4A and B**), in agreement with prior reports demonstrating that properly engineered catechol based polymers can be highly cytocompatible and bioactive. [33, 34, 35] Importantly, encapsulated cells within hPDA fueled fibrin constructs exhibited progressive spreading and matrix interaction, indicating that strong adhesion does not impede cellular remodeling. This balance between early mechanical stabilization and biological permissiveness is essential for meniscus repair, where intrinsic healing capacity is limited and zonally regulated. [46] The translational relevance of hPDA was most clearly demonstrated in the avascular bovine meniscus explant model, which closely recapitulates the biological, transport, and mechanical constraints of inner zone meniscal healing. [47, 48, 9] Recent studies have shown that meniscus cell migration and reparative responses are strongly zonal dependent and epigenetically constrained under inflammatory conditions, further limiting intrinsic healing in the avascular region. [46] In addition, transport of biological cues within meniscus fibrocartilage is highly restricted by matrix density, tissue layer, and molecular size, [49, 43, 5] underscoring the importance of early interface stabilization to sustain reparative signaling. Consistent with these insights, hPDA fueled bioglues promoted near continuous tissue bridging, dense collagen deposition, and pronounced fiber alignment across the repair interface (**Fig. 5B**), hallmarks of structurally mature healing tissue. In contrast, fibrin only and genipin crosslinked controls exhibited persistent interfacial gaps and disorganized collagen architecture (**Fig. 5B**).

These biological outcomes translated directly into restoration of mechanical function across length scales. Tensile pull out testing revealed large increases in interfacial stiffness (**Fig. 5C**) and strength (**Fig. 5D**), while nanoindentation based modulus mapping demonstrated elimination of low modulus interfacial regions (**Fig. 6A**) that typically act as stress concentrators. Restoration of interfacial stiffness toward native meniscus levels is particularly significant (**Fig. 6B**), as meniscus mechanical behavior, including tensile energy dissipation and load transfer, is highly dependent on fiber orientation, tissue layer, and hydration state. [50] Moreover, mechanical discontinuities within the meniscus have been shown to drive progressive matrix degeneration and cell pathology, even in compartment matched tissues, [51] emphasizing the clinical importance of restoring mechanical continuity rather than achieving nominal defect closure. Schematic illustration (**Fig. 7**) depicts the depolymerization driven transition from aggregated PDA to water soluble hPDA and the associated increase in functional group accessibility that underpins enhanced fibrin and tissue interactions.

**Figure 7:**
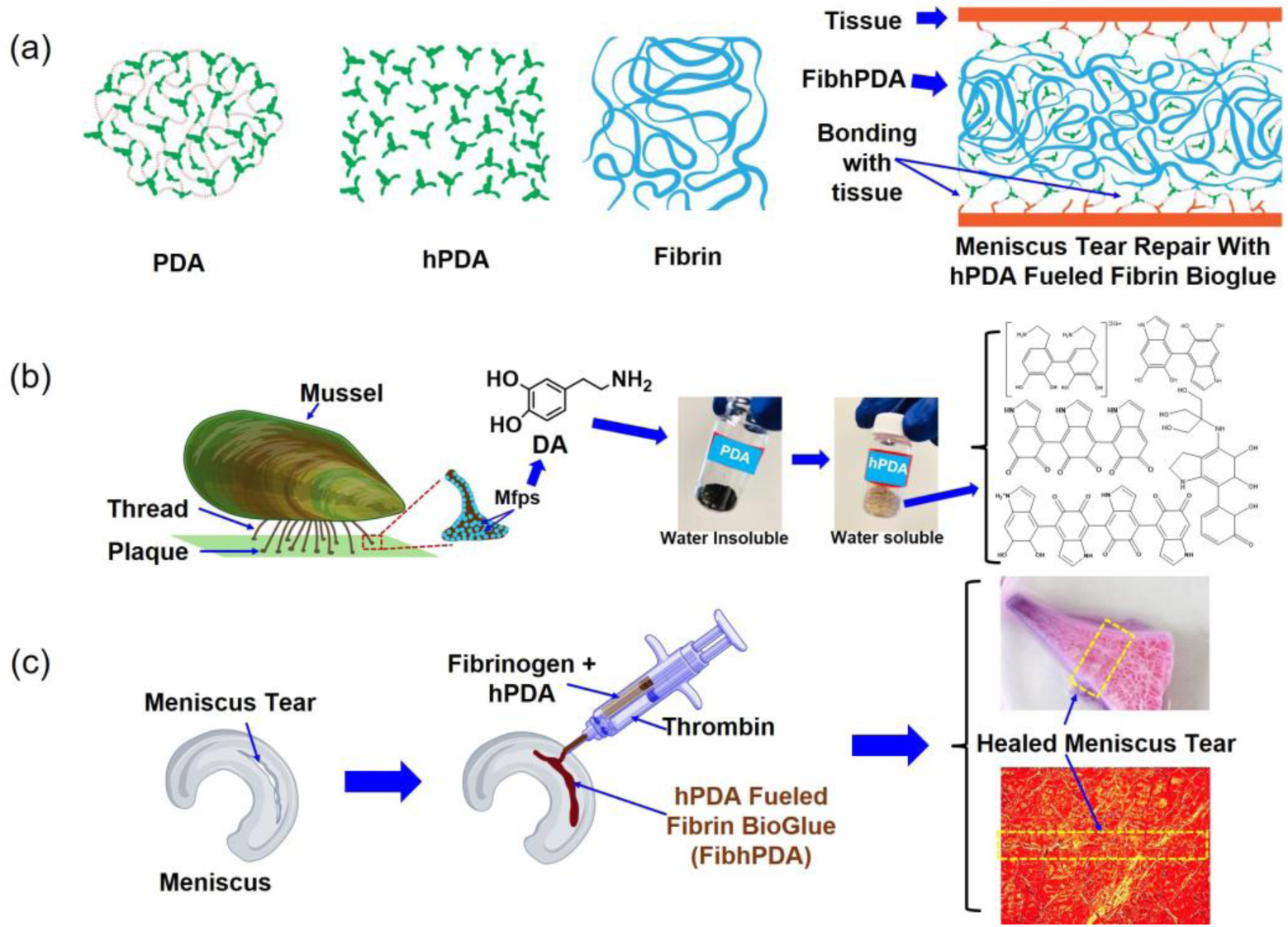
Schematic illustration of the design, formation mechanism, and translational application of hPDA fueled fibrin bioglue for avascular meniscus repair. (a) Top panels illustrate the molecular- and network-level concepts underlying the bioglue system. Conventional PDA consists of densely aggregated catechol rich polymer domains stabilized by strong intermolecular interactions. Controlled depolymerization and disruption of these interactions generate water-soluble hPDA monomers and oligomers with increased molecular mobility and enhanced accessibility of catechol, amine, and hydroxyl functional groups. These exposed functionalities enable multivalent interactions with fibrin and native extracellular matrix components, promoting strong interfacial bonding and improved mechanical stability when incorporated into a fibrin network. (b) Middle panel highlights the bioinspired origin of catechol-mediated adhesion from mussel foot proteins (Mfps) and illustrates the chemical transformation from insoluble PDA to hPDA, enabling integration into injectable and cell-compatible biomaterials. (c) Bottom panel depicts the translational application of the hPDA fueled fibrin bioglue (FibhPDA), which is delivered by injection into an avascular meniscus tear, resulting in defect filling, tissue bridging, and enhanced collagen organization at the repair interface, as demonstrated by representative gross and histological images of the treated region.

Despite the strong translational promise demonstrated in this study, several limitations should be considered. While the explant model provides a rigorous and clinically relevant assessment of avascular meniscus healing, it does not fully recapitulate the inflammatory environment, immune response, or complex multiaxial joint loading present in vivo. Future studies will therefore focus on evaluating hPDA fueled bioglues in large animal models under physiologic joint loading to assess long term durability, tissue integration, and remodeling. In addition, hPDA was evaluated as a heterogeneous population of oligomeric species. Although this approach reflects a practical and scalable synthesis strategy, future work aimed at refining molecular weight distribution and functional group density may allow more precise tuning of adhesive strength, degradation behavior, and biological interactions, building on recent advances in catechol based polymer design. [34, 35]

Collectively, these findings position hPDA as a hydrophilic, catechol based interfacial mediator that bridges materials innovation with biological and mechanical requirements for functional meniscus repair. Unlike adhesive systems that prioritize either adhesion or bioactivity, hPDA simultaneously enhances adhesion, mechanical performance, and biological integration using a synthesis strategy compatible with injectable and modular biomaterial platforms. Beyond meniscus repair, the material principles demonstrated here, interface dominated load transfer, degradation control, and cytocompatibility, are broadly relevant to other connective tissues characterized by limited vascularity and high mechanical demand, including tendon, ligament, cartilage, annulus fibrosus, and tendon to bone interfaces.

## 4. Conclusion

This study introduces hPDA as a transformative bioadhesive component that fundamentally advances the repair of avascular meniscus tears. By converting water-insoluble PDA into a stable, water-soluble hydrophilic form (hPDA), we unlocked its full interfacial reactivity and enabled seamless integration within fibrin-based bioglues. hPDA incorporation significantly enhanced adhesive stiffness, strength, viscoelastic integrity, and degradation control, outperforming fibrin alone and rivaling genipin-crosslinked systems while preserving excellent cytocompatibility. Importantly, hPDA fueled bioglues promoted superior biological and mechanical healing in a bovine avascular meniscus explant model, characterized by continuous tissue integration, aligned collagen remodeling, and restoration of interfacial mechanical properties approaching native meniscus tissue across macro- and micro-scales. Multimodal mechanical analyses revealed that hPDA is the primary contributor to improved load transfer and stiffness continuity, with genipin providing complementary but limited reinforcement. The hPDA fueled bioglue strategy offers a clinically relevant, injectable, and mechanically robust solution with strong translational potential to improve long-term outcomes following meniscal injury and reduce the progression toward osteoarthritis. Beyond meniscus repair, the unique combination of strong adhesion, biological compatibility, and mechanical reinforcement conferred by hPDA suggests broad applicability across a wide range of connective tissues that suffer from poor intrinsic healing. Tissues such as tendon, ligament, cartilage, and annulus fibrosus share common challenges, including limited vascularity, high mechanical demand, and frequent failure at repair interfaces. The ability of hPDA to enhance interfacial load transfer, stabilize healing under dynamic loading, and support matrix remodeling positions it as a promising adhesive and interfacial mediator for these tissues.

## 5. Materials and Methods

### 5.1. Materials

Dopamine hydrochloride (F8630, Sigma-Aldrich, St. Louis, MO, USA), hydrochloric acid (258148, Sigma-Aldrich, St. Louis, MO, USA), sodium hydroxide (30620, Sigma-Aldrich, St. Louis, MO, USA), and Tris base (11814273001, Roche, Indianapolis, IN, USA) were used for hPDA synthesis.Thrombin (605157, Sigma-Aldrich, St. Louis, MO, USA), genipin (G4796, Sigma-Aldrich, St. Louis, MO, USA), fibrinogen (F8630, Sigma-Aldrich, St. Louis, MO, USA), and human bone marrow-derived mesenchymal stem cells(hBMSCs,SCC034, Sigma-Aldrich, St. Louis, MO, USA) were used for cytotoxicity, degradation, lap-shear, and tensile tests. Thrombin (605157, Sigma-Aldrich, St. Louis, MO, USA), genipin (G4796, Sigma-Aldrich, St. Louis, MO, USA), fibrinogen (F8630, Sigma-Aldrich, St. Louis, MO, USA), hBMSCs (SCC034, Sigma-Aldrich, St. Louis, MO, USA), connective tissue growth factor (CTGF, human, SRP4702, Sigma-Aldrich, St. Louis, MO, USA), and transforming growth factor-*β*3 (TGF-*β*3, human, H8791, Sigma-Aldrich, St. Louis, MO, USA) were used. Additionally, 1% ITS+1 solution (I2521, Sigma-Aldrich, St. Louis, MO, USA), sodium pyruvate (P2256, Sigma-Aldrich, St. Louis, MO, USA), L-ascorbic acid 2-phosphate (A8960, Sigma-Aldrich, St. Louis, MO, USA), L-proline(A10199.14, Sigma-Aldrich, St. Louis, MO, USA), and dexamethasone(D4902, Sigma-Aldrich, St. Louis, MO, USA) were used in the explant culture medium.

### 5.2. Hydrophilic Polydopamine (hPDA) Synthesis

Water-soluble hPDA was synthesized from water-insoluble PDA particles using a modified base-dissolution and recrystallization approach. Briefly, dopamine hydrochloride (5 mg/mL) was first dissolved in TRIS/HCl buffer (pH 8.5) and stirred overnight at room temperature in a glass beaker. This reaction led to the formation of insoluble PDA particles dispersed throughout the solution. Following polymerization, PDA particles were isolated by centrifugation, washed sequentially with deionized water and absolute ethanol, and air-dried. The dried PDA was then dissolved in a sodium hydroxide (NaOH) solution to break down the insoluble PDA structures. Subsequently, excess ethanol was added to the alkaline solution to induce crystallization of hPDA as fine precipitates. These precipitated hPDA particles were thoroughly washed with 100% ethanol to remove residual impurities and then air-dried to yield the final hPDA powder. The synthesized hPDA was used for the formulation of bioglues.

### 5.3. hPDA Characterization

The synthesized hPDA revealed its excellent water solubility, which represents a significant improvement over DA. The physicochemical and morphological properties of the synthesized hPDA were evaluated using Ultravio-let–Visible (UV–Vis) spectroscopy, Fourier Transform Infrared Spectroscopy (FTIR), Mass Spectrometry (MS), and Scanning Electron Microscopy (SEM). These characterization techniques were employed to confirm the successful transformation of conventional, water-insoluble PDA into a water-soluble hPDA form and to assess changes in optical properties, chemical structure, and particle morphology. UV–Vis spectroscopic analysis of DA and hPDA was performed over the 190–400 nm wavelength range to identify spectral features indicative of hPDA formation. FTIR spectra were acquired using a Spectrum 100 FT-IR spectrometer (Perkin Elmer, France) in universal attenuated total reflection (UATR) mode with a Diamond/KRS-5 crystal. Both PDA and hPDA samples were analyzed across a wavenumber range of 400 to 4000 *cm*^−1^ to identify key functional group vibrations and structural differences between the two materials. The surface morphology of hPDA particles was observed using a Hitachi S-4700 scanning electron microscope. Samples were first lyophilized to ensure complete dryness, mounted on aluminum stubs using double-sided conductive carbon tape, and sputter-coated with a gold layer (20–30 nm thick) to improve electrical conductivity. Mass spectrometry analysis was performed using a Sciex 5500+ triple quadrupole instrument to investigate the molecular composition and assess structural transitions from DA to PDA and subsequently to hPDA. Samples were introduced using positive ion electrospray ionization (ES^+^) at a flow rate of 5*µ*L/min. Instrument settings included a spray voltage of 5.5 kV, curtain gas at 20 psi, ion source gas 1 at 20 psi, ion source gas 2 at 0 psi, and a source temperature of 0°C. This analysis enabled detection of mass spectral signatures that reflect changes in molecular weight and fragmentation patterns.

### 5.4. Rheological Characterization

The rheological properties of the bioglues were evaluated using a TA Instruments AR2000ex rheometer (TA Instruments, New Castle, DE, USA) equipped with a 40 mm parallel plate geometry and a fixed gap of 500 *µ*m. Rheological measurements were conducted at 37°C unless otherwise specified. Shear-dependent flow behavior was determined in controlled shear rate mode by measuring viscosity and shear stress over a shear rate range of 0.1*s*^−1^-1000*s*^−1^. The linear viscoelastic region (LVR) was identified by a strain sweep from 0.1% to 1000% strain at 1 Hz, monitoring storage (*G*′) and loss (*G*^′′^) moduli. Frequency-dependent viscoelastic behavior was evaluated within the LVR at a constant strain of 4% and at frequencies ranging from 0.1 to 100 Hz. Three replicates (n = 3) were tested for each group.

### 5.5. Degradation Kinetics of the hPDA Fueled Bioglue

For the in vitro degradation study, four bioadhesive formulations were evaluated: fibrin (Fib), hPDA fueled fibrin (FibhPDA), genipin crosslinked fibrin (FibGen), and genipin crosslinked hPDA fueled fibrin (FibGenhPDA). Our previous findings demonstrated that the incorporation of genipin into fibrin bioglue significantly slows degradation and enhances adhesive performance [45]. In this study, genipin-containing groups served both as a benchmark to compare against hPDA and to assess whether combining genipin with hPDA yields any synergistic benefit. A previously optimized genipin concentration of 2.5 mg/mL was used for all crosslinked formulations [45]. Each bioadhesive formulation was supplemented with Alexa Fluor® 488 (10 *µ*L/mL) to enable fluorescence-based monitoring of degradation dynamics over time. Equal volumes of each formulation (50 *µ*L; N = 6 per group) were cast into 24-well plates and incubated in PBS at 37 °C. Fluorescence and corresponding bright-field images were collected at predetermined time points (2, 3, 4, 7, and 14 days) using a Maestro™ in vivo fluorescence imaging system (Cambridge Research & Instrumentation, Inc., Woburn, MA, USA). Degradation kinetics were quantified by measuring the residual bioglue area over time using ImageJ (NIH).

### 5.6. Compression Test

The compressive moduli of the bioglues were evaluated using a CellScale UniVert uniaxial mechanical testing system (CellScale, Waterloo, Canada) equipped with a 50 N load cell. Square-shaped hydrogel specimens (approximately 5 mm × 5 mm × 2 mm) were prepared from the same four formulations used for the lap-shear testing: (i) fibrin alone (Fib), (ii) hPDA fueled fibrin (FibhPDA), (iii) genipin-crosslinked fibrin (FibGen), and (iv) genipin-crosslinked hPDA fueled fibrin (FibGenhPDA). Compression tests were conducted at a crosshead speed of 3.9 mm/min at room temperature, and samples were compressed to 90% of their initial thickness. The compressive moduli were determined from the linear portion of the stress vs strain plots within the elastic deformation region. A minimum of six (*n* ≥ 6) were tested for each group.

### 5.7. Lap-Shear Test

The adhesive strength of the hPDA fueled bioglue was evaluated using our established lap-shear test procedure on bovine meniscus tissue [48, 47, 45]. Briefly, menisci were harvested from skeletally mature bovine knee joints, and the inner-third zone was sectioned into wedge-shaped strips with a thickness of 2–4 mm and approximate dimensions of 25 mm × 10 mm. Four hydrogel formulations (i) fibrin alone (Fib), (ii) hPDA fueled fibrin (FibhPDA), (iii) genipin-crosslinked fibrin (FibGen), and (iv) genipin-crosslinked hPDA fueled fibrin (FibGenhPDA) were tested. For each test, 20 *µ*L of the bioglue was applied to the contact region between two meniscus strips, creating a defined overlap area of approximately 5 mm × 10 mm. The samples were gently pressed together and incubated at 37°C in a humidified chamber for 30 minutes to ensure complete gelation and bonding. Following curing, lap-shear tests were conducted using the CellScale UniVert uniaxial mechanical testing system (Waterloo, Canada) equipped with a 10 N load cell. The samples were mounted such that tensile load was applied parallel to the bonded interface at a constant displacement rate of 0.1 mm/s (equivalent to 5 mm/min). The maximum force at failure (*F_max_*) was recorded, and the adhesive shear strength (*σ*) was calculated using the formula:

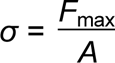

where A represents the bonded area (mm^2^). The lap-shear modulus was also calculated from the linear slope of the stress–strain curve within the elastic deformation region. A minimum of six replicates (*n* ≥ 6) were tested for each group.

### 5.8. Cytotoxicity

The cytocompatibility of synthesized hydrophilic hPDA was evaluated using a live/dead cell viability assay in both two-dimensional (2D) and three-dimensional (3D) cultures of human bone marrow-derived mesenchymal stem cells (hBMSCs) for seven days. For the 2D assay, hBMSCs were cultured in DMEM alone or in DMEM supplemented with hPDA or DA. Cell viability was assessed to determine any potential cytotoxic effects of hPDA. In the 3D model, hPDA fueled fibrin (FibhPDA) bioglue was prepared by adding hPDA with fibrinogen solution followed by mixing with thrombin and hBM-SCs to form a cell-laden construct. The final concentrations of the individual components in the bioglue formulation were 100 mg fibrinogen, 100 U thrombin, 6 mg hPDA, and 6 × 10^6^ cells per mL bioglue. A control fibrin gel was prepared using the same protocol but without hPDA.

### 5.9. Explant Model for Avascular Meniscus Healing

To evaluate the role of hPDA in promoting tissue integration and repair, our well-established bovine meniscus explant model was used to mimic avascular meniscal injuries [47, 48]. Medial and lateral menisci were collected from skeletally mature bovine knee joints (Animal Technologies, Inc., Tyler, TX 75702). The inner one-third region of each meniscus was isolated, and radial sections were made to obtain wedge-shaped explants approximately 5 mm in thickness, yielding 5–6 explants per meniscus. A controlled, full-thickness longitudinal incision was created in the center of each explant using a sterile surgical blade. The defect was filled with 50 *µ*L of the bioglue from the four hydrogel formulations: (i) fibrin alone (Fib), (ii) hPDA fueled fibrin (FibhPDA), (iii) genipin-crosslinked fibrin (FibGen), and (iii) genipin-crosslinked hPDA fueled fibrin (FibGenhPDA). All hydrogel formulations were supplemented with connective tissue growth factor (CTGF, 100 ng/mL) and transforming growth factor-*β*3 (TGF*β*3) encapsulated PLGA microspheres (10 mg PLGA *µ*S/mL bioglue). Treated explants were cultured atop a confluent (80 to 90%) monolayer of passage 2 to 3 human bone marrow-derived mesenchymal stem cells (hBMSCs). A 1:1 mixture of fibrogenic and chondrogenic differentiation media was used throughout the explant culture period, following protocols established in our previous studies [47, 48]. The fibrogenic medium contained 50 *µ*g/mL ascorbic acid, while the chondrogenic medium was supplemented with 1% ITS+1, 100 *µ*g/mL sodium pyruvate, 50 *µ*g/mL L-ascorbic acid 2-phosphate, 40 *µ*g/mL L-proline, and 0.1 *µ*M dexamethasone. After a 6-week culture period, explants were harvested and analyzed for healing of the avascular tear using histological evaluation, biochemical assays, and multi-scale mechanical testing.

### 5.10. Tensile Testing for Tissue Integration

To assess the effectiveness of hPDA feuled bioglues in promoting meniscal tissue integration, uniaxial tensile (pull-out) tests were conducted following 6 weeks of explant culture. Each unfixed explant was secured in custom tensile grips at room temperature, and a preload of 0.02 N was applied prior to testing. The samples were then stretched at a constant rate of 10% strain per minute until failure. Throughout the testing process, specimens were kept hydrated with frequent spray of phosphate-buffered saline (PBS). Mechanical parameters including maximum force, tensile strength, and tensile modulus were derived from the stress vs strain plot derived. All mechanical tests were performed using a UniVert testing platform (CellScale, Waterloo, ON, Canada).

### 5.11. Modulus Mapping

Elastic modulus mapping was performed to characterize the micro-mechanical properties of the repaired tissue within the avascular region of the meniscus explants, following our previously established procedures [47, 48]. Briefly, unfixed and unstained tissue sections were mounted on a high-precision motorized X–Y stage of a PIUMA™ nanoindenter (Optics11, Amsterdam, The Netherlands) equipped with a 10 *µ*m spherical probe. A maximum indentation force of 10 mN was applied at 20 *µ*m intervals across the healed interface to capture spatial variations in the effective elastic modulus (*E_Eff_*).For each experimental group, five regions were mapped, yielding 100–150 indentation points per region. The resulting modulus measurements were plotted as spatial maps based on X–Y coordinates, and average *E_Eff_* values were computed over 100 *µ*m wide segments spanning the repair interface, consistent with our previously published methodology [47, 48].

### 5.12. Statistical Analysis

All quantitative data were analyzed using one-way analysis of variance (ANOVA) followed by Tukey’s HSD post-hoc test to determine statistical significance. A significance level of *α* = 0.05 was used for all comparisons. Sample sizes for each quantitative dataset were determined by a priori power analysis using one-way ANOVA with *α* = 0.05, power = 0.8, and an effect size of 1.50.

## Supporting Information

Additional supporting information can be found online in the Supporting Information section.

## Acknowledgements

This material is based upon work supported by the National Science Foundation under the Engineering Research Initiation (ERI) Award No. 2347673 (to S. Tarafder). Any opinions, findings, and conclusions or recommendations expressed in this material are those of the authors and do not necessarily reflect the views of the National Science Foundation.The authors also acknowledge the South Dakota State University Campus Mass Spectrometry Facility for use of the Agilent GC–MS/MS instrumentation used in this work. The GC–MS/MS was acquired through funding from the U.S. Department of Agriculture through a National Institute of Food and Agriculture Equipment Grants Program (Award No. 2023-70410-41209).

## CRediT authorship contribution statement

**Hasan Rafsan Jani:** Writing – review & editing, Writing – original draft, Investigation, Methodology, Data curation, Visualization. **Maya A. Jeremias:** Writing – review & editing, Investigation, Methodology, Data curation. **Aminah T. Sarowar:** Writing – review & editing, Investigation, Methodology, Data curation. **M. Nurul Islam:** Writing – review & editing, Data curation, Formal analysis. **Chang H. Lee:** Writing – review & editing, Data curation, Formal analysis, Supervision. **Solaiman Tarafder:** Writing – review & editing, Writing – original draft, Conceptualization, Methodology, Data curation, Formal analysis, Validation, Visualization, Resources, Supervision, Project administration.

## Data Availability Statement

Data will be made available on request.

---

**Figure S1:**
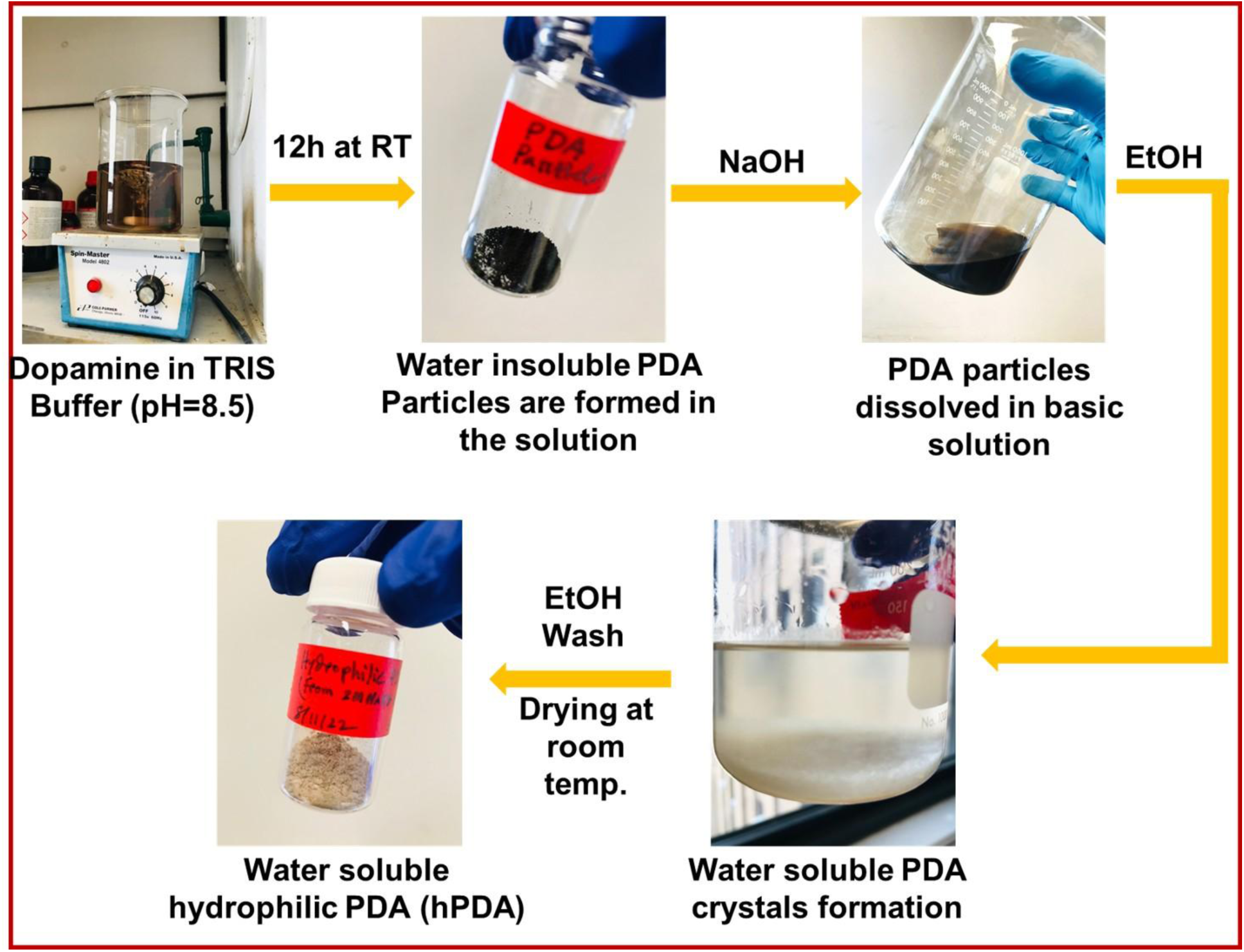
Synthesis workflow for hydrophilic polydopamine (hPDA). Dopamine was oxidized in TRIS buffer (pH 8.5) at room temperature for 12 h to form water-insoluble PDA particulates. The crude PDA was dissolved under basic conditions (NaOH) and treated with ethanol to induce crystallization of water-soluble PDA intermediates. Following ethanol washing and ambient drying, a water-soluble hydrophilic PDA (hPDA) powder was obtained. Photographs illustrate each step from initial dopamine solution to final hPDA product; RT, room temperature; EtOH, ethanol.

**Figure S2:**
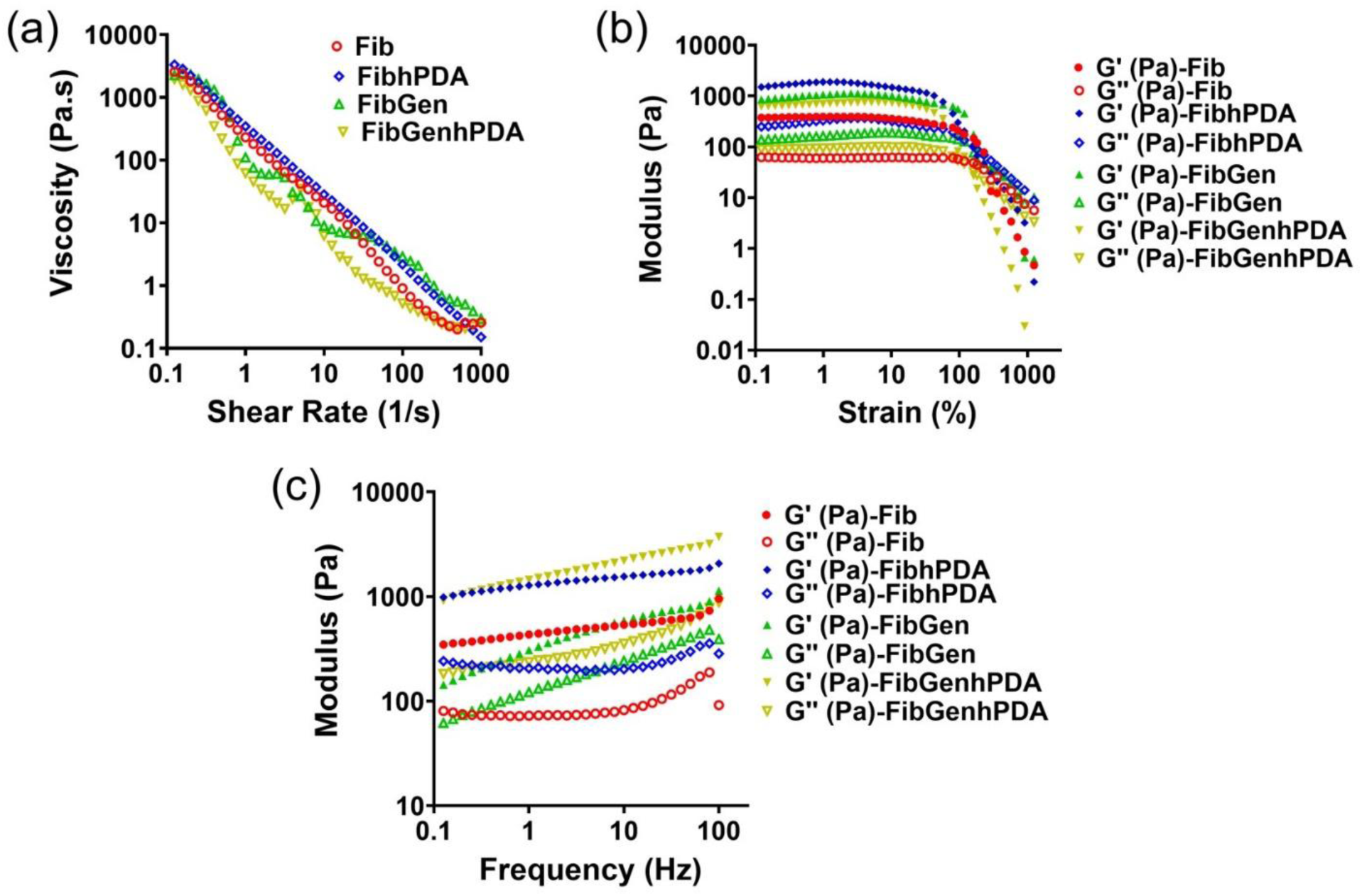
Rheological characterization of fibrin and hPDA based bioadhesives. (A) Viscosity versus shear rate showing pronounced shear-thinning behavior for all formulations (Fib, FibGen, FibhPDA, FibGenhPDA) across 0.1–1000s^−1^ shear rate. (B) Strain sweep at 0.1-1000 strain(%) identifying the linear viscoelastic region (LVR); storage modulus G′ exceeds loss modulus G′′ at low strains for all gels, with higher moduli observed for hPDA and genipin-containing formulations until yield. (C) Frequency sweep within the LVR (4% strain, 0.1–100 Hz) demonstrating gel-like behavior (G′ > G′′) and increased moduli for FibhPDA and FibGenhPDA relative to Fib and FibGen. All characterization was done at 37 °C using a 40 mm parallel-plate geometry with 500 μm gap.

## References

[1] J. A. Koski, C. Ibarra, S. A. Rodeo, R. F. Warren, Meniscal injury and repair: clinical status, Orthopedic Clinics 31 (3) (2000) 419–435.

[2] E. A. Makris, P. Hadidi, K. A. Athanasiou, The knee meniscus: structure–function, pathophysiology, current repair techniques, and prospects for regeneration, Biomaterials 32 (30) (2011) 7411–7431.

[3] H. S. Cheung, Distribution of type i, ii, iii and v in the pepsin solubilized collagens in bovine menisci, Connective tissue research 16 (4) (1987) 343–356.

[4] J. N. Wasserburger, C. L. Shultz, D. A. Hankins, L. Korcek, D. F. Martin, A. Amendola, D. L. Richter, R. C. Schenck, G. P. Treme, Long-term national trends of arthroscopic meniscal repair and debridement, The American Journal of Sports Medicine 49 (6) (2021) 1530–1537.

[5] E. M. Marigi, S. E. Till, J. N. Wasserburger, A. K. Reinholz, A. J. Krych, M. J. Stuart, Inside-out approach to meniscus repair: Still the gold standard?, Current Reviews in Musculoskeletal Medicine 15 (4) (2022) 244–251.

[6] B. M. Baker, A. O. Gee, N. P. Sheth, G. R. Huffman, B. J. Sennett, T. P. Schaer, R. L. Mauck, Meniscus tissue engineering on the nanoscale– from basic principles to clinical application, The journal of knee surgery 22 (01) (2009) 45–59.

[7] F. R. Noyes, S. D. Barber-Westin, Repair of complex and avascular meniscal tears and meniscal transplantation, JBJS 92 (4) (2010) 1012–1029.

[8] H. Kwon, W. E. Brown, C. A. Lee, D. Wang, N. Paschos, J. C. Hu, K. A. Athanasiou, Surgical and tissue engineering strategies for articular cartilage and meniscus repair, Nature Reviews Rheumatology 15 (9) (2019) 550–570.

[9] R. M. Irwin, M. Brown, M. F. Koff, C. H. Lee, E. Lemmon, H. J. Jeong, S. P. Simmonds, J. L. Robinson, A. M. Seitz, P. Tanska, et al., Generating new meniscus therapies via recent breakthroughs in development, model systems, and clinical diagnostics, Journal of Orthopaedic Research® 43 (6) (2025) 1073–1089.

[10] R. Papalia, A. Del Buono, L. Osti, V. Denaro, N. Maffulli, Meniscectomy as a risk factor for knee osteoarthritis: a systematic review, British medical bulletin 99 (1) (2011) 89–106.

[11] A. L. McNulty, F. Guilak, Mechanobiology of the meniscus, Journal of biomechanics 48 (8) (2015) 1469–1478.

[12] S. R. Lee, J. G. Kim, S. W. Nam, The tips and pitfalls of meniscus allograft transplantation, Knee surgery & related research 24 (3) (2012) 137.

[13] M. M. Pillai, J. Gopinathan, R. Senthil Kumar, G. Sathish Kumar, S. Shanthakumari, K. S. Sahanand, A. Bhattacharyya, R. Selvakumar, Tissue engineering of human knee meniscus using functionalized and reinforced silk-polyvinyl alcohol composite three-dimensional scaffolds: understanding the in vitro and in vivo behavior, Journal of Biomedical Materials Research Part A 106 (6) (2018) 1722–1731.

[14] M. Koch, F. P. Achatz, S. Lang, C. G. Pfeifer, G. Pattappa, R. Kujat, M. Nerlich, P. Angele, J. Zellner, Tissue engineering of large full-size meniscus defects by a polyurethane scaffold: accelerated regeneration by mesenchymal stromal cells, Stem cells international 2018 (1) (2018) 8207071.

[15] G. Desando, G. Giavaresi, C. Cavallo, I. Bartolotti, F. Sartoni, N. Nicoli Aldini, L. Martini, A. Parrilli, E. Mariani, M. Fini, et al., Autologous bone marrow concentrate in a sheep model of osteoarthritis: new perspectives for cartilage and meniscus repair, Tissue Engineering Part C: Methods 22 (6) (2016) 608–619.

[16] Y.-F. Zhou, D. Zhang, W.-T. Yan, K. Lian, Z.-Z. Zhang, Meniscus regeneration with multipotent stromal cell therapies, Frontiers in Bioengineering and Biotechnology 10 (2022) 796408.

[17] Y.-H. An, H. D. Kim, K. Kim, S.-H. Lee, H.-G. Yim, B.-G. Kim, N. S. Hwang, et al., Enzyme-mediated tissue adhesive hydrogels for meniscus repair, International journal of biological macromolecules 110 (2018) 479–487.

[18] A. Bochyńska, G. Hannink, R. Verhoeven, D. Grijpma, P. Buma, Evaluation of novel biodegradable three-armed-and hyper-branched tissue adhesives in a meniscus explant model, Journal of Biomedical Materials Research Part A 105 (5) (2017) 1405–1411.

[19] J. Zellner, C. D. Taeger, M. Schaffer, J. C. Roldan, M. Loibl, M. B. Mueller, A. Berner, W. Krutsch, M. K. Huber, R. Kujat, et al., Are applied growth factors able to mimic the positive effects of mesenchymal stem cells on the regeneration of meniscus in the avascular zone?, BioMed Research International 2014 (1) (2014) 537686.

[20] F. Liu, H. Xu, H. Huang, A novel kartogenin-platelet-rich plasma gel enhances chondrogenesis of bone marrow mesenchymal stem cells in vitro and promotes wounded meniscus healing in vivo, Stem cell research & therapy 10 (1) (2019) 201.

[21] M. Horie, M. D. Driscoll, H. W. Sampson, I. Sekiya, C. T. Caroom, D. J. Prockop, D. B. Thomas, Implantation of allogenic synovial stem cells promotes meniscal regeneration in a rabbit meniscal defect model, JBJS 94 (8) (2012) 701–712.

[22] M. Horie, I. Sekiya, T. Muneta, S. Ichinose, K. Matsumoto, H. Saito, T. Murakami, E. Kobayashi, Intra-articular injected synovial stem cells differentiate into meniscal cells directly and promote meniscal regeneration without mobilization to distant organs in rat massive meniscal defect, Stem cells 27 (4) (2009) 878–887.

[23] Y. Nakagawa, T. Muneta, S. Kondo, M. Mizuno, K. Takakuda, S. Ichinose, T. Tabuchi, H. Koga, K. Tsuji, I. Sekiya, Synovial mesenchymal stem cells promote healing after meniscal repair in microminipigs, Osteoarthritis and Cartilage 23 (6) (2015) 1007–1017.

[24] F. Duygulu, M. Demirel, G. Atalan, F. Kaymaz, Y. Kocabey, T. Dulgeroglu, H. Candemir, Effects of intra-articular administration of autologous bone marrow aspirate on healing of full-thickness meniscal tear: an experimental study on sheep, Acta orthopaedica et traumatologica turcica 46 (1) (2012) 61–67.

[25] C. T. Jayasuriya, J. Twomey-Kozak, J. Newberry, S. Desai, P. Feltman, J. R. Franco, N. Li, R. Terek, M. G. Ehrlich, B. D. Owens, Human cartilage-derived progenitors resist terminal differentiation and require cxcr4 activation to successfully bridge meniscus tissue tears, Stem Cells 37 (1) (2019) 102–114.

[26] S.-J. Heo, K. H. Song, S. Thakur, L. M. Miller, X. Cao, A. P. Peredo, B. N. Seiber, F. Qu, T. P. Driscoll, V. B. Shenoy, et al., Nuclear softening expedites interstitial cell migration in fibrous networks and dense connective tissues, Science advances 6 (25) (2020) eaax5083.

[27] A. Bochyńska, T. Van Tienen, G. Hannink, P. Buma, D. W. Grijpma, Development of biodegradable hyper-branched tissue adhesives for the repair of meniscus tears, Acta biomaterialia 32 (2016) 1–9.

[28] L. Han, M. Wang, P. Li, D. Gan, L. Yan, J. Xu, K. Wang, L. Fang, C. W. Chan, H. Zhang, et al., Mussel-inspired tissue-adhesive hydrogel based on the polydopamine–chondroitin sulfate complex for growth-factor-free cartilage regeneration, ACS applied materials & interfaces 10 (33) (2018) 28015–28026.

[29] H. Lee, B. P. Lee, P. B. Messersmith, A reversible wet/dry adhesive inspired by mussels and geckos, Nature 448 (7151) (2007) 338–341.

[30] Y. Fu, L. Yang, J. Zhang, J. Hu, G. Duan, X. Liu, Y. Li, Z. Gu, Poly-dopamine antibacterial materials. mater horizons 8: 1618–1633 (2021).

[31] A. Jin, Y. Wang, K. Lin, L. Jiang, Nanoparticles modified by poly-dopamine: Working as “drug” carriers, Bioactive materials 5 (3) (2020) 522–541.

[32] L. Ai, Y. Wang, G. Tao, P. Zhao, A. Umar, P. Wang, H. He, Polydopamine-based surface modification of zno nanoparticles on sericin/polyvinyl alcohol composite film for antibacterial application, Molecules 24 (3) (2019) 503.

[33] J. H. Ryu, P. B. Messersmith, H. Lee, Polydopamine surface chemistry: a decade of discovery, ACS applied materials & interfaces 10 (9) (2018) 7523–7540.

[34] N. Yin, Z. Zhang, Y. Ge, Y. Zhao, Z. Gu, Y. Yang, L. Mao, Z. Wei, J. Liu, J. Shi, et al., Polydopamine-based nanomedicines for efficient antiviral and secondary injury protection therapy, Science Advances 9 (24) (2023) eadf4098.

[35] Y. Liu, D. He, Y. Cheng, L. Li, Z. Lu, R. Liang, Y. Fan, Y. Qiao, S. Chou, A heterostructure coupling of bioinspired, adhesive poly-dopamine, and porous prussian blue nanocubics as cathode for high-performance sodium-ion battery, Small 16 (11) (2020) 1906946.

[36] J. Liebscher, R. Mrówczyński, H. A. Scheidt, C. Filip, N. D. Hădade, R. Turcu, A. Bende, S. Beck, Structure of polydopamine: a never-ending story?, Langmuir 29 (33) (2013) 10539–10548.

[37] X. Chen, W. Yang, J. Zhang, L. Zhang, H. Shen, D. Shi, Alkalinity triggered the degradation of polydopamine nanoparticles, Polymer Bulletin 78 (8) (2021) 4439–4452.

[38] H. Jia, Z. Su, W. Long, Y. Liu, X. Chang, H. Zhang, G. Ding, Y. Feng, D. Cai, Z. Zou, Metabonomics combined with uplc-ms chemical profile for discovery of antidepressant ingredients of a traditional chinese medicines formula, chaihu-shu-gan-san, Evidence-Based Complementary and Alternative Medicine 2013 (1) (2013) 487158.

[39] Q. Lyu, N. Hsueh, C. L. Chai, Unravelling the polydopamine mystery: is the end in sight?, Polymer Chemistry 10 (42) (2019) 5771–5777.

[40] N. F. Della Vecchia, A. Luchini, A. Napolitano, G. D’Errico, G. Vitiello, N. Szekely, M. d’Ischia, L. Paduano, Tris buffer modulates polydopamine growth, aggregation, and paramagnetic properties, Langmuir 30 (32) (2014) 9811–9818.

[41] K. Patel, N. Singh, J. Yadav, J. M. Nayak, S. K. Sahoo, J. Lata, D. Chand, S. Kumar, R. Kumar, Polydopamine films change their physicochemical and antimicrobial properties with a change in reaction conditions, Physical Chemistry Chemical Physics 20 (8) (2018) 5744–5755.

[42] H. Fan, X. Yu, Y. Liu, Z. Shi, H. Liu, Z. Nie, D. Wu, Z. Jin, Folic acid– polydopamine nanofibers show enhanced ordered-stacking via *π*–*π* interactions, Soft Matter 11 (23) (2015) 4621–4629.

[43] A. Morejon, G. Schwartz, T. M. Best, F. Travascio, A. R. Jackson, Effect of molecular weight and tissue layer on solute partitioning in the knee meniscus, Osteoarthritis and cartilage open 5 (2) (2023) 100360.

[44] J. H. Ryu, S. Hong, H. Lee, Bio-inspired adhesive catechol-conjugated chitosan for biomedical applications: A mini review, Acta biomaterialia 27 (2015) 101–115.

[45] S. Tarafder, J. Ghataure, D. Langford, R. Brooke, R. Kim, S. L. Eyen, J. Bensadoun, J. T. Felix, J. L. Cook, C. H. Lee, Advanced bioactive glue tethering lubricin/prg4 to promote integrated healing of avascular meniscus tears, Bioactive Materials 28 (2023) 61–73.

[46] Y. Zhang, E. Y. Zhang, C. Cheung, Y. Heo, B.-I. Tumenbayar, S.-H. Lee, Y. Bae, S. C. Heo, Epigenetic dynamics in meniscus cell migration and its zonal dependency in response to inflammatory conditions, APL bioengineering 9 (1) (2025).

[47] S. Tarafder, J. Gulko, K. H. Sim, J. Yang, J. L. Cook, C. H. Lee, Engineered healing of avascular meniscus tears by stem cell recruitment, Scientific reports 8 (1) (2018) 8150.

[48] S. Tarafder, J. Gulko, D. Kim, K. H. Sim, S. Gutman, J. Yang, J. L. Cook, C. H. Lee, Effect of dose and release rate of ctgf and tgf*β*3 on avascular meniscus healing, Journal of Orthopaedic Research® 37 (7) (2019) 1555–1562.

[49] G. Schwartz, S. Rana, A. R. Jackson, C. Leñero, T. M. Best, D. Kouroupis, F. Travascio, Human mesenchymal stem/stromal cell-derived extracellular vesicle transport in meniscus fibrocartilage, Journal of Orthopaedic Research® 43 (2) (2025) 457–465.

[50] A. Morejon, P. L. Dalbo, T. M. Best, A. R. Jackson, F. Travascio, Tensile energy dissipation and mechanical properties of the knee meniscus: relationship with fiber orientation, tissue layer, and water content, Frontiers in bioengineering and biotechnology 11 (2023) 1205512.

[51] S. G. Lopez, J. Dankert, J. J. Butler, J. G. Kennedy, L. J. Bonassar, R. M. Irwin, Medial osteochondral defect drives matrix and cell pathology in compartment-matched meniscus, Journal of Orthopaedic Research® 43 (11) (2025) 2031–2042.

